# Mitochondrial and metabolic plasticity distinguish stem cells from progenitors in the male germline

**DOI:** 10.64898/2026.10.08.757768

**Authors:** Katerina B. Damyanova, Joshua J. Fisher, Siddharth Acharya, Brett Nixon, Tessa Lord

**Affiliations:** Centre for Reproductive Science, College of Engineering, Science and Environment, University of Newcastle, Callaghan, NSW, 2308, Australia; Reproductive and Family Health Research Program, Hunter Medical Research Institute, New Lambton Heights, NSW 2305, Australia; School of Medicine and Public Health, College of Health, Medicine and Wellbeing, University of Newcastle, Callaghan, NSW, Australia; School of Biomedical Sciences and Pharmacy, College of Health, Medicine and Wellbeing, University of Newcastle, Callaghan, NSW, Australia

**Keywords:** spermatogonial stem cells, metabolic plasticity, cristae organisation, oxidative phosphorylation, glycolysis, redox balance

## Abstract

Metabolism and mitochondrial remodelling are increasingly recognised as key regulators of stem cell identity, governing the balance between self-renewal and differentiation. Here we compared mitochondrial and metabolic properties of stem and progenitor cells from the male germline using transmission electron microscopy, quantitative proteomics and extracellular flux analysis. While the stem cell-to-progenitor transition was accompanied by increased mitochondrial abundance and cristae density, our analyses unveiled a remarkable and unexpected capacity for spermatogonial stem cells (SSCs) to engage oxidative phosphorylation. When oxygen availability was unrestricted, SSCs displayed markedly greater maximal and spare respiratory capacity than progenitors, despite comparable glycolytic flux. SSCs were also enriched for glutathione-dependent antioxidant enzymes and maintained lower basal mitochondrial superoxide levels than progenitors. Collectively, these findings challenge the view that SSCs are predominantly glycolytic and instead identify metabolic plasticity, marked by latent oxidative capacity and enhanced redox control, as a hallmark of the stem cell state.

## INTRODUCTION

Metabolism and metabolic plasticity are now recognised as active regulators of stem cell identity, functioning as dynamic developmental switches that guide cells between self-renewal and differentiation (reviewed in (Lord and Nixon, 2020)). Stem cell metabolic states are shaped not only by the conditions of their niche, particularly oxygen availability, but also by the maturity and functional capacity of their mitochondria. Understanding how microenvironmental cues and mitochondrial plasticity interact to support stem cell maintenance versus lineage commitment is therefore indispensable for advancing *in vitro* stem cell applications. In the male germline, spermatogonial stem cells (SSCs) generate spermatozoa in the adult testis, and SSC-based therapies have a broad range of applications, from clinical infertility treatments to wildlife conservation (reviewed in (Damyanova et al., 2024)). Although recent work has begun to define the SSC niche and the link between glycolytic metabolism and self-renewal (Bernstein et al., 2023; Helsel et al., 2017a; Kanatsu-Shinohara et al., 2016), comparatively little is known about the ultrastructural or functional characteristics of SSC mitochondria. This gap in knowledge contributes to persistent challenges in expanding SSCs in culture while preserving long-term regenerative capacity (Helsel et al., 2017a).

To sustain lifelong spermatogenesis, SSCs must balance self-renewal with the production of progenitor spermatogonia primed for differentiation (Lord and Oatley, 2017). A key regulator of SSC fate is the surrounding microenvironment, which provides essential signalling factors and maintains physiological conditions that shape stem cell behaviour (Oatley and Brinster, 2012). SSCs reside in a relatively hypoxic microenvironment at the basement membrane of seminiferous tubules, whereas progenitor and differentiating spermatogonia are located closer to the interstitial space, where they are exposed to higher oxygen levels (Bernstein et al., 2023; Lord and Nixon, 2020). This spatial organisation underpins a broader principle observed across stem cell systems, where undifferentiated cells preferentially rely on glycolysis to meet their bioenergetic demands, while differentiation is accompanied by a metabolic shift toward oxidative phosphorylation (OXPHOS) (Ito and Suda, 2014). Consistent with this model, independent studies show that stimulating glycolysis through overexpression of MYC/MYCN (Kanatsu-Shinohara et al., 2016) or upregulation of hypoxia-responsive pathways (Bernstein et al., 2023; Helsel et al., 2017a), promotes SSCs self-renewal.

Because mitochondrial structure and function govern metabolic states, defining SSC-specific mitochondrial features is essential for understanding how metabolic programs support stem cell maintenance versus lineage commitment. However, isolating pure SSC populations has historically been technically challenging, owing to their close morphological and transcriptional similarity to undifferentiated progenitors (Chan et al., 2014; Helsel et al., 2017b; Xi et al., 2022). As a result, no studies have characterised the ultrastructure or functional properties of SSC mitochondria, nor their capacity for metabolic plasticity under different conditions. While transcriptomic analyses suggest that mitochondrial protein expression changes during the transition from SSC to progenitor (Guo et al., 2017; Hermann et al., 2018; Lord and Nixon, 2020), our recent multiomic analyses have demonstrated poor correlation between transcript and protein abundance in spermatogonia (Skerrett-Byrne et al., 2026), consistent with observations in the testis more broadly (Jiang et al., 2020). Together these findings highlight a critical gap in our understanding of the mitochondrial properties that delineate self-renewing SSCs from their differentiation-primed progenitor counterparts.

In contrast to the limited understanding of SSC-specific mitochondrial features, the metabolic and mitochondrial changes accompanying the process of spermatogonial differentiation have been more clearly defined. Following a retinoic acid pulse in the testis, undifferentiated progenitor spermatogonia make the differentiating transition, preparing them for meiosis and ultimately their transformation into spermatozoa. During this transition, spermatogonia exhibit an upregulation of OXPHOS and an increase in mitochondrial fission and fusion (Zhang et al., 2024). Disruption of either fission (Zhang et al., 2024) or fusion (Chen et al., 2020a) blocks differentiation, highlighting the requirement for a metabolic shift in this process. These mitochondrial changes parallel those observed in other stem cell lineages, where differentiation is often accompanied by broader mitochondrial remodelling, including expansion of mitochondrial DNA (mtDNA) copy number and redistribution of mitochondria throughout the cytoplasm to support increased reliance on mitochondrial energy production (Facucho-Oliveira et al., 2007).

Beyond the role of metabolism in dictating stem cell identity, the by-products of metabolism, reactive oxygen species (ROS), are also key regulators of cell fate. ROS are not solely damaging by-products of mitochondrial respiration but also function as important signalling molecules that influence proliferation and differentiation. In several stem cell systems, tightly regulated ROS levels promote differentiation, whereas excessive oxidative stress can compromise stem cell maintenance and genomic stability (Bigarella et al., 2014; Holmström and Finkel, 2014). In the male germline, ROS signalling plays a particularly important role, with NOX1-derived ROS being reported to promote SSC self-renewal through p38/JNK-dependent pathways (Morimoto et al., 2013). Additionally, SSCs possess unusually robust antioxidant and DNA-repair systems, including high base-excision repair activity mediated by OGG1, to protect against ROS-induced damage (Mori et al., 2021). Despite these insights, virtually nothing is known about how ROS levels or antioxidant capacity differ between SSCs and their undifferentiated progenitor counterparts, or how redox signalling integrates with the metabolic and mitochondrial changes that accompany the onset of differentiation. Consistent with a role for redox regulation in this process, inhibition of mitochondrial respiration, mitochondrial dynamics, or ROS production blocks spermatogonial differentiation (Chen et al., 2020b; Varuzhanyan et al., 2019; Zhang et al., 2024). Thus, defining redox balance across spermatogonial subpopulations is essential, particularly for *in vitro* applications where maintaining genomic integrity is crucial for producing healthy sperm capable of generating viable offspring.

Despite advances in understanding SSC biology, the long-term maintenance of SSCs *in vitro* remains a significant challenge. Current culture systems allow for SSC expansion, yet stem cell function declines after several months, limiting their utility in clinical and research settings (Helsel et al., 2017a; Kubota et al., 2009; Schmidt et al., 2010). This limitation is particularly problematic given the rarity of SSCs in the testis (Tagelenbosch and de Rooij, 1993), necessitating prolonged culture periods to obtain sufficient cell numbers for downstream applications. Whether this loss of function reflects underlying disruptions to SSC metabolism, mitochondrial dynamics, or redox balance remains unknown, but preliminary transcriptomic comparisons suggest that metabolic dysregulation may contribute to culture-induced dysfunction (De Oliveira et al., 2022). Identifying the precise metabolic requirements of SSCs is therefore critical for developing culture conditions that preserve long-term stem cell function.

In this study, we provide the first comprehensive comparison of mitochondrial and metabolic features in stem cell and progenitor populations within the male germ line. By integrating mtDNA quantification, assessment of mitochondrial localisation, morphology and cristae organisation, metabolic activity measurements (oxygen consumption rate (OCR) and proton efflux rate (PER)), and a proteomic analysis of pathways governing metabolism and redox balance, paired with a kinetic metabolic model, we define how these populations differ in both metabolic state and metabolic potential. Together these approaches reveal a striking degree of mitochondrial and metabolic plasticity that distinguishes the stem cell pool from progenitors. Although SSC mitochondria display cristae architecture consistent with lower electron-transport efficiency, SSCs retain an exceptional capacity to upregulate OXPHOS when shifted out of their hypoxic niche, displaying an energetic reserve that far exceeds that of progenitors. These findings uncover previously unrecognised metabolic flexibility in SSCs and highlight mitochondrial state as a key determinant of stem cell identity, an insight with important implications for the optimisation of long-term SSC culture and future stem cell-based applications.

## MATERIALS AND METHODS

### Animals

All procedures involving animal use were approved by the University of Newcastle Animal Care and Ethics Committee (ACEC, approval numbers A-2019-907, A-2024-410 and A-2024-412). The *Id4-eGfp* mouse line (Chan et al., 2014) was a gift from Dr Jon Oatley.

### Cell sorting

PD6–8 testes were dissociated into a single-cell suspension by incubating for 5 min at 37°C in 0.25% Trypsin-EDTA containing 7mg DNase I (Sigma Aldrich), followed by gentle pipetting. The cell suspension was kept on ice in Dulbecco’s Phosphate Buffered Saline (DPBS, Sigma Aldrich) containing 10% Fetal Bovine Serum (FBS, Cell Sera Australia, NSW, Australia), hereafter referred to as DPBSS. ID4-eGFP^Bright^ and ID4-eGFP^Dim^ populations of spermatogonia were sorted using a BD FACS Aria II using previously described gating parameters (Helsel et al., 2017b).

### MitoTracker Immunofluorescence

To examine the subcellular localisation of mitochondria in both SSCs and progenitors we used Thy-1 cell surface antigen (THY1+) testis cell populations isolated from postnatal day (PD) 6– 8 testes via magnetic activated cell sorting (MACS). Testes were dissociated into a single-cell suspension via 5 min incubation at 37°C in 0.25% Trypsin-EDTA containing 7mg DNase I (Sigma Aldrich) and cells were labelled with a biotin-conjugated Thy1 antibody (Invitrogen; #13-0902-82) and separated using anti-biotin microbeads (Miltenyi Biotech, North Rhine-Westphalia, Germany; #130-104-253) following the manufacturer’s instructions. The THY1+ cell population encompasses both ID4-eGFP^Bright^ and ID4-eGFP^Dim^ cells, allowing for a direct comparison through immunocytochemistry. Isolated cells were incubated with 100 nM MitoTracker® Deep Red (Invitrogen; #M22426) for 30 min at 37°C, washed twice in Hanks’ Balanced Salt Solution (HBSS, Gibco; #14025092) and fixed in 4% paraformaldehyde for 15 min at 37°C. After two washes in 0.1 M glycine in PBS, cells were settled onto Poly-L-Lysine coated cover slips in a humidified chamber overnight at 4°C. The next day, cells were permeabilised with 0.2% Triton X-100 for 10 min at room temperature, washed twice with PBS containing 0.1% Tween-20 (PBST) and counterstained with DAPI (1:1000) for 5 min. Coverslips were washed in PBST and mounted onto slides using Mowiol. Imaging was performed on a Zeiss LSM 900 confocal microscope using a 63x oil objective. Z-stacks spanning the full cellular volume were acquired, and images are presented as sum-intensity projections.

### Mitochondrial DNA (mtDNA) copy number assessment

To compare mitochondrial content between SSCs and progenitor spermatogonia, relative mitochondrial DNA (mtDNA) copy number was determined using quantitative polymerase chain reaction (qPCR) as previously described (Facucho-Oliveira et al., 2007). Following FACS, ID4-eGFP^Bright^ and ID4-eGFP^Dim^ cells were snap frozen and total DNA (genomic and mitochondrial) was extracted using the Monarch Genomic DNA Purification Kit (New England Biolabs; #T3010) following the manufacturer’s instructions. DNA concentration and purity were assessed using a NanoDrop® (ThermoFisher).

qPCR reactions (10 µL) contained 2.0 ng DNA, 1× GoTaq® Master Mix (Promega) and 0.33 µM of forward and reverse primers (Invitrogen Life Technologies). Each sample was run in triplicate. A 366 bp product of the nuclear *Gapdh* gene was amplified with *Gapdh*-F (5’-GGGAAGCCCATCACCATCTTC-3’) and *Gapdh*-R (5’-AGAGGGGCCATCCACAGTCT-3’); and 211 bp product of the *tRNA-Tyr* and *mt-Co1* genes of the mtDNA, amplified with *tRNA-Tyr/mt-Co1*-F (5’-CAGTCTAATGCTTACTCAGC-3’) and *tRNA-Tyr/mt-Co1*-R (5’-GGGCAGTTACGATAACATTG-3’). This mtDNA locus was selected because it is highly conserved, present in a single copy per mitochondrial genome (Bibb et al., 1981), and avoids regions with known nuclear mitochondrial pseudogene interference (Mishmar et al., 2004), making it a reliable target for mtDNA copy-number quantification. Amplification was performed on a LightCycler® 96 Real-Time PCR system with the following conditions; *tRNA-Tyr/mt-Co1*: 95°C for 10 min, followed by 50 cycles of 95°C for 10 sec, 53°C for 15 sec, and 72°C for 15 sec; *Gapdh*: 95°C for 10 min, followed by 50 cycles of 95°C for 10 sec, 60°C for 15 sec, and 72°C for 23 sec.

Fluorescence was measured at the final step of each cycle using the FAM/SYBR channel. Relative mtDNA copy number was calculated using the comparative Ct (ΔCt) method, normalising the mitochondrial target (*tRNA-Tyr/mt-Co1*) to the nuclear reference gene *Gapdh*. Technical triplicates were averaged, and relative mtDNA copy number was expressed as 2 × 2^(Ct(Gapdh) − Ct(tRNA-Tyr/mt-Co1))^, with the factor of 2 accounting for the two copies of Gapdh per diploid nuclear genome.

### Mitochondrial Membrane Potential (MMP)

PD6–8 testes were dissociated into a single-cell suspension via 5 min incubation at 37°C in 0.25% Trypsin-EDTA containing 7mg DNase I (Sigma Aldrich). Cells were then incubated with 10 nM Tetramethylrhodamine (TMRM, Invitrogen; #I34361) for 30 minutes at 37°C to assess mitochondrial membrane potential (MMP), followed by washing in DPBSS. As a depolarisation control, cells were treated with 10 µM of the mitochondrial uncoupler CCCP for 1h post-TMRM staining to confirm an MMP-dependent reduction in TMRM fluorescence. Cells were again washed in DPBSS and fluorescence was measured using a BD FACSCanto II Flow Cytometer.

### MitoSOX Red (MSR)

To assess mitochondrial superoxide production, PD7 testes were pooled from 2–4 animals per replicate and dissociated into single-cell suspensions as described above. Following dissociation, cells from n = 4 independent biological replicates were stained with 5 μM MitoSOX Red (MSR; Invitrogen; #M36008) and 50 nM SYTOX Deep Red (Invitrogen; #S11380) for 30 minutes at 37°C protected from light, washed in DPBSS and analysed immediately using a BD FACSCanto II flow cytometer. Events were gated sequentially to exclude debris and SYTOX^+^ non-viable cells (**Supplementary Fig. S1**), followed by identification of ID4-eGFP^+^ undifferentiated spermatogonia and separation of ID4-eGFP^Bright^ (SSCs) and ID4-eGFP^Dim^ (progenitor) subpopulations. MitoSOX Red fluorescence was quantified as the proportion of MSR^+^ events within total ID4-eGFP^Bright^ and ID4-eGFP^Dim^ populations. Unstained and single-colour controls were used for gate setting and compensation. Final numerical values were generated from a complete FlowJo v11 reanalysis using a consistent gating strategy applied across all samples.

### Electron Microscopy

For transmission electron microscopy (TEM) analysis, ID4-eGFP^Bright^ and ID4-eGFP^Dim^ cell populations were isolated from PD7 testes (n = 14) using FACS, as described above. Cells were pooled into SSC and progenitor populations and fixed for 1 h at room temperature in TEM fixative containing 2.5% glutaraldehyde, 4% paraformaldehyde, and 100 mM sodium cacodylate. Following two washes in 100 mM cacodylate buffer, cells were embedded in 2% low–melting-point agarose (Sigma Aldrich; #A9415). Agarose pellets were then post-fixed overnight at 4°C in fresh TEM fixative, with fixative changes at 1 and 2 h. After three 5 min washes in cacodylate buffer, samples were post-fixed in 1% osmium tetroxide (ProSciTech; #C011) for 2 h at 4°C and washed four times in Milli-Q water. Samples were dehydrated through a graded acetone series (30%, 50%, 70%, 90%, and four changes of 100% acetone; 15 min each), infiltrated with an acetone–resin mixture, embedded in EMbed 812 resin (EMS; #14120), and polymerised at 60°C for 24 h. Ultrathin sections (100 nm) were cut using a diamond knife on a Leica EM UC7 ultramicrotome, stretched with chloroform to eliminate compression and mounted on 200-mesh TEM nickel grids, and imaged using a JEM F-200 CFEG multipurpose S/TEM at 20,000x magnification. Given that SSCs have a reported diameter of approximately 11–12 µm (Martin and Seandel, 2013), multiple sections spaced at least 12 µm apart were collected from each resin block to generate technical replicates.

### Quantitative analysis of mitochondrial morphology

Representative images were imported into Amira 3D (Thermo Fisher Scientific, USA), where sigma and non-local means filters were applied to enhance mitochondrial membrane contrast prior to cristae segmentation. Manual segmentation was performed on mitochondria from SSCs (ID4-eGFP^Bright^; n = 77 mitochondria) and progenitor spermatogonia (ID4-eGFP^Dim^; n = 80 mitochondria) by tracing only clearly discernible mitochondrial inner and outer membranes using ImageJ v1.54 (NIH). Mitochondrial ultrastructure was assessed using previously described parameters, including mitochondrial surface area, major (length) and minor (width) axes, and elongation (length:width ratio) (Choi et al., 2020; Lam et al., 2021).

Given the inherent limitations of single-section (two-dimensional) TEM for resolving three-dimensional cristae topology, tubular and globular cristae could not be reliably distinguished in all instances. In contrast, lamellar cristae exhibited well-defined membrane outlines and junctions that could be consistently identified across sections. Accordingly, quantitative cristae analyses focused on lamellar cristae parameters, including surface area, density (total cristae-to-mitochondrial surface area ratio), major and minor axes, elongation ratio, and cristae junction width, consistent with approaches used in recent mitochondrial ultrastructure studies (Acharya et al., 2026a) (**Supplementary Fig. S2**). Classification of cristae into lamellar, tubular, and globular categories was therefore used for descriptive purposes only. Additional morphological descriptors included mitochondrial perimeter, circularity, and solidity, which were used to assess membrane complexity and dynamic remodelling associated with mitochondrial fusion–fission states (Picard et al., 2013). In this context, increased perimeter and reduced circularity are indicative of elongated or irregular mitochondrial morphology, while lower solidity reflects branching or angular structures and higher solidity is consistent with mitochondrial fusion (Westrate et al., 2014).

### Seahorse analyses

Oxygen consumption rate (OCR) and extracellular acidification rate (ECAR) were measured using an XF96 Extracellular Flux Analyzer (Seahorse Bioscience, North Billerica, MA, USA). ID4-eGFP^Bright^ and ID4-eGFP^Dim^ cells were seeded at 5 × 10⁴ cells per well in XF96 cell culture microplates in mouse serum-free medium (mSFM) supplemented with glial cell-derived neurotrophic factor (GDNF; 20 ng/mL; R&D Systems, MN, USA) and fibroblast growth factor 2 (FGF2; 1 ng/mL; Sigma-Aldrich) and incubated overnight at 37°C in 10% O_2_ and 5% CO_2._ On the day of the assay, culture medium was replaced with Seahorse XF DMEM (Agilent Technologies; #103575-100) supplemented with 10mM glucose, 1mM sodium pyruvate, and 2mM L-glutamine. Cells were incubated for 1 h at 37°C and 21% O_2_ in a CO_2_-free incubator prior to analysis, and fresh assay medium was added immediately before measurements.

For assessment of mitochondrial respiration (Mito Stress Test), OCR was measured under basal conditions and following sequential injections of oligomycin (1.5 μM; Sigma-Aldrich; #O4876), FCCP (1 μM; Abcam; #ab120081), and a combination of rotenone (0.5 μM; Sigma-Aldrich; #R8875) and antimycin A (0.5 μM; Sigma-Aldrich; #A8674). Basal respiration was calculated from baseline OCR measurements prior to inhibitor injection. ATP-linked respiration was defined as the decrease in OCR following oligomycin treatment, reflecting oxygen consumption coupled to ATP synthesis. Maximal respiration was determined following FCCP injection, which uncouples oxidative phosphorylation and drives the electron transport chain to its maximal capacity. Spare respiratory capacity was calculated as the difference between maximal and basal respiration.

Glycolytic activity was assessed using the Seahorse Glycolytic Rate Assay (Agilent Technologies; #103344-100). PER was derived from ECAR using the buffer factor of the assay medium (2.5 mmol H⁺/pH/L) and the instrument volume constants, and was measured under basal conditions and following sequential injections of rotenone/antimycin A (0.5 μM each) to inhibit mitochondrial respiration and drive compensatory glycolysis, and 2-deoxy-D-glucose (2-DG; 50 mM) to inhibit glycolysis. As mitochondrial CO_2_ also contributes to extracellular acidification, glycolytic PER (glycoPER) was calculated as glycoPER = PER − (CO_2_ contribution factor × mitochondrial OCR), using the default CO_2_ contribution factor of 0.61 and taking mitochondrial OCR as total OCR minus the non-mitochondrial rate measured following rotenone/antimycin A injection; this correction therefore applies only to basal measurements. Basal glycolysis was taken as glycoPER at the final basal measurement, compensatory glycolysis as the maximal glycoPER following rotenone/antimycin A injection, and post-2-DG acidification as glycoPER following 2-DG injection. The proportion of proton efflux attributable to glycolysis was calculated as basal glycoPER divided by basal PER. The relative contribution of mitochondrial respiration and glycolysis was assessed as the ratio of basal mitochondrial OCR to basal glycoPER (mitoOCR:glycoPER).

To verify cell seeding density, assay wells were imaged immediately prior to running the assay using the EVOS™ M5000 Imaging System (Thermo Fisher Scientific, Waltham, MA, USA) (**Supplementary Fig. S3**). Seahorse-derived parameters were normalised to post-attachment cell counts obtained from representative imaging fields. Technical replicate wells were averaged to generate a single value per biological replicate prior to statistical analysis.

### Daprodustat treatment of cultured spermatogonia

Primary cultures of undifferentiated spermatogonia, established from ID4-eGFP^+^ spermatogonia isolated from PD6–8 testes, were maintained on mitotically inactivated STO feeder cells in mSFM supplemented with GDNF (20 ng/mL) and FGF2 (1 ng/mL) at 37°C, 10% O_2_ and 5% CO_2_, as previously described (Bernstein et al., 2023). Five independent cultures (n = 5; passages 5–8) were used as biological replicates. Spermatogonia were dislodged from the feeder layer by gentle pipetting in HBSS, dissociated in 0.25% Trypsin-EDTA for 5 min at 37°C, quenched with 10% FBS, pelleted at 600 × g for 7 min and counted. Cells were seeded at 5 × 10^4^ cells per well in XF96 cell culture microplates in 80 µL mSFM with GDNF and FGF2, containing either 125 µM Daprodustat (MedChemExpress; #HY-17608) or an equivalent volume of DMSO vehicle (0.5% v/v), with two to three technical replicate wells per condition. Plates were then incubated for 24 h at 37°C, 10% O_2_ and 5% CO_2_. The Glycolytic Rate Assay was then performed as described above, without Daprodustat in the assay medium.

### Proteomics

To define metabolic and mitochondrial pathway differences between SSCs and progenitor spermatogonia, we performed *in silico* analysis on a previously published proteomics dataset from our research group (Skerrett-Byrne et al., 2026). This dataset captures spermatogonia isolated from PD6-8 *Id4-eGfp* mice, where ID4-eGFP^Bright^ and ID4-eGFP^Dim^ populations were isolated using equivalent methods to those described above (Skerrett-Byrne et al., 2026). Fold change and significance (t-test) comparative testing between sample groups was carried out within the Proteome Discoverer 2.5 suite. The peptide list was exported from Proteome Discoverer 2.5 as an Excel file and further refined to include only those with a quantitative value in all replicates of at least one of the sample groups. The dataset was then imported into Perseus for normalisation and scaling. Proteins of interest were identified by filtering against two published datasets of metabolic (Corcoran et al., 2017) and mitochondrial proteins (Rath et al., 2020), supplemented with a UniProt search using the key words ‘metabolism’, ‘oxidative phosphorylation’, ‘glycolysis’ and ‘mitochondria’ (see **Supplementary Table S1** for protein lists). For pathway-focused visualisation, curated proteins were subsequently grouped into functional categories and displayed as individual heat maps representing proteins involved in mitochondrial fragmentation and structural remodelling, core glycolytic enzymes and transcription factor regulators, electron transport chain (ETC) complexes I–V, and regulators of ROS production and redox balance.

### Quantitative systems metabolism (QSM) modelling

To assess potential metabolic capacities based on the proteomic dataset, protein abundance values were analysed using Quantitative Systems Metabolism (QSM™) modelling (Doppelganger Biosystems GmbH, Oldenburg, Germany) (Berndt et al., 2021; Berndt et al., 2015). This computational framework integrates quantitative proteomic data with curated metabolic network models to estimate the relative capacity of metabolic pathways and substrate utilisation. Protein intensity values for SSCs and progenitor spermatogonia were mapped onto central carbon metabolic pathways, including glycolysis, the tricarboxylic acid cycle, fatty acid oxidation, and lactate metabolism. The resulting model outputs were used to compare predicted ATP production capacity and substrate utilisation potential between the two populations.

Model outputs corresponding to the postprandial metabolic state were used for comparative analyses, as germ cells within the seminiferous epithelium exist in a nutrient-replete environment supported by continuous metabolite exchange with Sertoli cells (Boussouar and Benahmed, 2004).

### Statistical Analysis

Statistical analyses were performed using GraphPad Prism v11.1.0, which was also used to generate heat maps and volcano plots for proteomic data visualisation. Quantitative TEM datasets were screened for outliers using the ROUT method (Q = 1%), and identified outliers were excluded. Data normality was assessed using the Shapiro–Wilk test. As TEM parameters did not meet normality assumptions in one or both populations, all were compared using the non-parametric Mann–Whitney U test. Seahorse-derived glycolytic and OXPHOS parameters, mitochondrial DNA copy number, mitochondrial membrane potential, QSM-modelled metabolic capacities and the proportion of MSR⁺ events were compared between SSCs and progenitor spermatogonia, and Glycolytic Rate Assay parameters between vehicle– and Daprodustat-treated cultured spermatogonia, using paired two-tailed t-tests. For all paired analyses, technical replicates were averaged to give a single value per biological replicate before testing. Statistical significance was defined as p < 0.05. Data are presented as mean ± standard deviation (SD) unless otherwise stated.

## RESULTS

### Assessing mitochondrial abundance and membrane potential in stem and progenitor populations of the germline lineage

To investigate whether mitochondrial abundance and membrane potential differ between SSCs and progenitor spermatogonia, we employed the *Id4-eGfp* transgenic mouse model, which enables prospective isolation of SSCs (ID4-eGFP^Bright^) and progenitor spermatogonia (ID4-eGFP^Dim^) by fluorescence-activated cell sorting (FACS) (Chan et al., 2014; Helsel et al., 2017b). Mitochondrial abundance was assessed by MitoTracker® Deep Red labelling and mtDNA copy number quantification (**Fig. 1**). Representative confocal images of MitoTracker® Deep Red-labelled THY1⁺ spermatogonia revealed visibly stronger mitochondrial-associated fluorescence in progenitor and differentiating spermatogonia (ID4-eGFP^Dim/negative^) relative to SSCs (ID4-eGFP^Bright^), consistent with greater mitochondrial abundance in cells that have progressed beyond the stem cell state (**Fig. 1A**). Subsequent quantification of relative mtDNA copy number revealed a significant increase in progenitor spermatogonia compared with SSCs (7.76 ± 2.33 vs 6.21 ± 1.50; *p* = 0.026; **Fig. 1B**), confirming greater mitochondrial abundance in the progenitor population.

**Figure 1.**
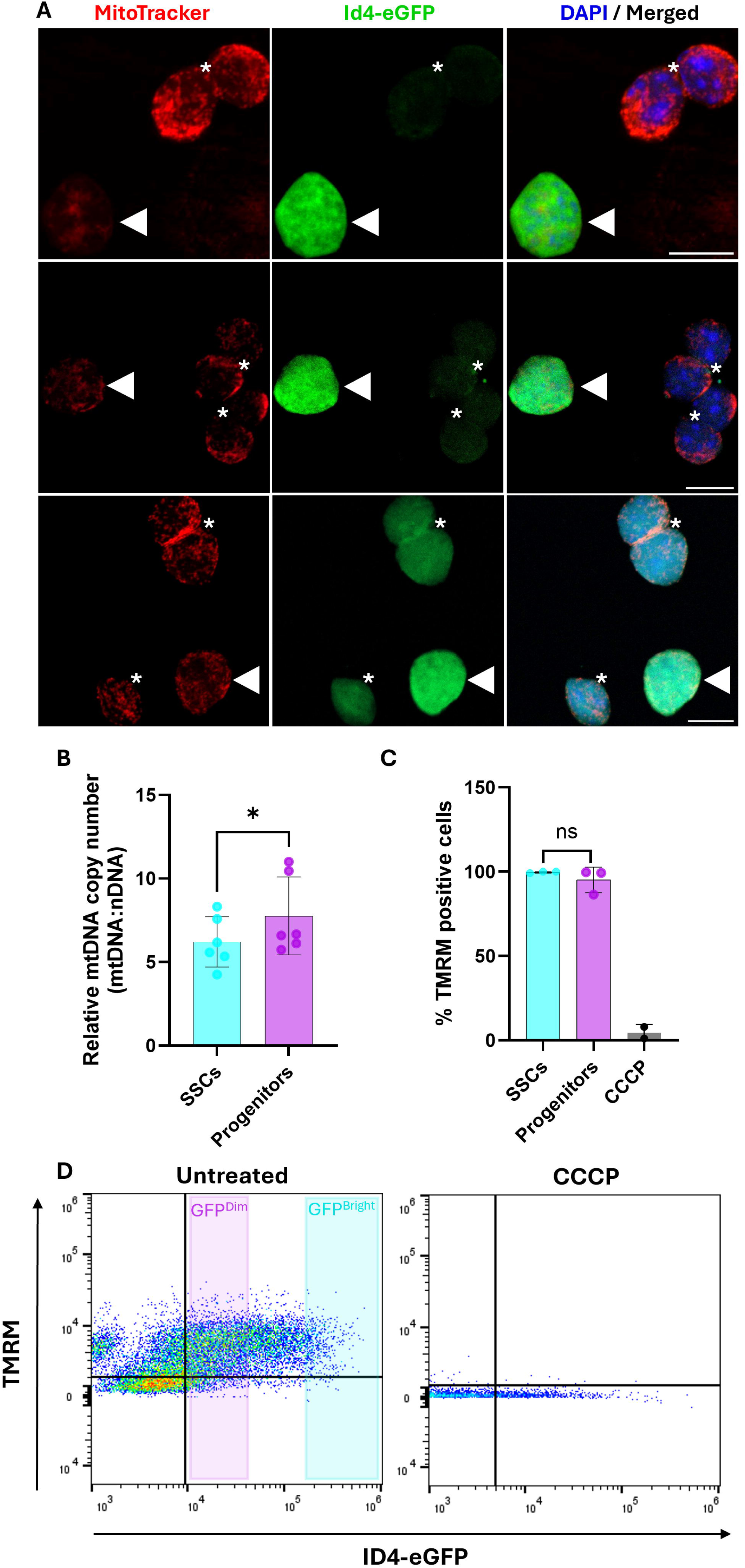
Mitochondrial abundance and membrane potential in spermatogonial stem cells (SSCs) and progenitor spermatogonia. (**A**) Representative confocal images of THY1⁺ undifferentiated spermatogonia stained with MitoTracker® Deep Red, showing higher overall mitochondrial fluorescence in GFP^Dim/negative^ progenitor and differentiating spermatogonia (asterisk) compared with GFP^Bright^ SSCs (arrowhead). Nuclei are counterstained with DAPI. Images represent sum-intensity projections of z-stacks. Scale bar = 10 µm **(B)** Relative mitochondrial DNA (mtDNA) copy number (mtDNA:nDNA ratio), determined by qPCR of the mitochondrial *tRNA-Tyr/mt-Co1* locus normalised to the nuclear *Gapdh* gene, was significantly higher in progenitor spermatogonia than in SSCs (n = 6; paired two-tailed t-test; p = 0.0256). **(C)** Mitochondrial membrane potential, assessed by TMRM staining and flow cytometry, was not significantly different between SSCs and progenitor spermatogonia (n = 3; paired two-tailed t-test; p = 0.3795). **(D)** Representative flow cytometry scatter plots showing TMRM fluorescence against ID4-eGFP signal in untreated cells (left) and cells treated with the protonophore CCCP (right), which dissipates the mitochondrial membrane potential.

To determine whether this difference in abundance was accompanied by differences in mitochondrial polarisation, membrane potential was assessed by TMRM staining and flow cytometry. The specificity of TMRM labelling was confirmed by treatment with the protonophore CCCP, which abolished membrane potential in both populations (**Fig. 1C, D**). No significant difference in membrane potential was detected between SSCs (99.72 ± 0.48) and progenitor spermatogonia (95.13 ± 7.58; *p* = 0.38; **Fig. 1C**). This indicates that while progenitor spermatogonia contain more mitochondria, individual mitochondria maintain comparable polarisation across both populations.

### Mitochondrial cristae architecture differs between SSCs and progenitor spermatogonia

To investigate whether mitochondrial organisation differs between SSCs and progenitor spermatogonia, we performed TEM on FACS-isolated ID4-eGFP^Bright^ SSCs (**Fig. 2A**) and ID4-eGFP^Dim^ progenitor spermatogonia (**Fig. 2B**), followed by quantitative ultrastructural analysis (**Fig. 2C-F**). Quantitative analysis of mitochondrial size and shape revealed no significant differences in mitochondrial surface area (*p* = 0.654; **Fig. 2C**), length (*p* = 0.591; **Fig. 2D**), width (*p* = 0.483; **Fig. 2E**), or elongation (length:width ratio; *p* = 0.603; **Fig. 2F**) between SSCs and progenitors, indicating comparable gross mitochondrial morphology across cell states.

**Figure 2.**
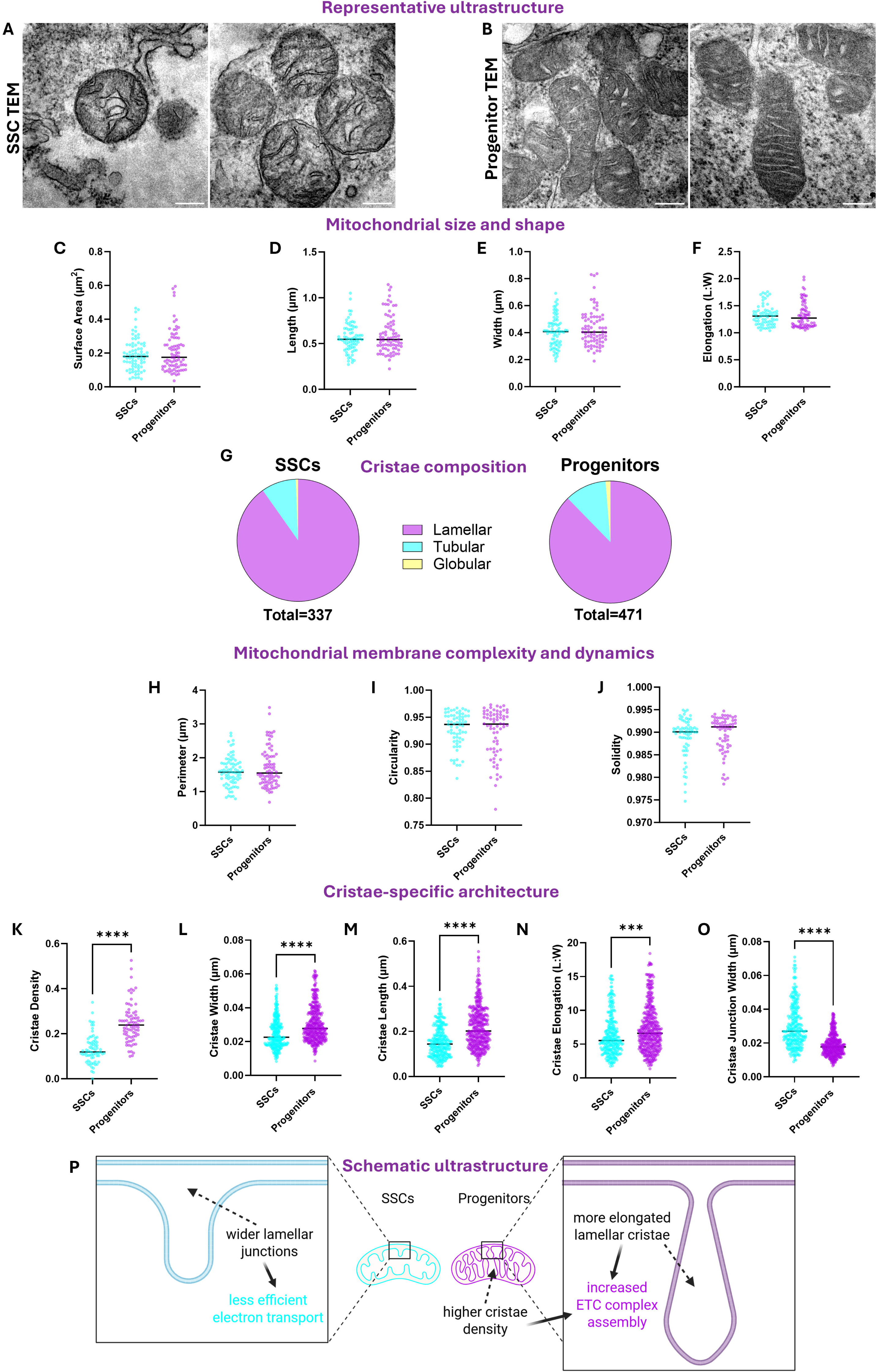
TEM analysis of mitochondrial ultrastructure and cristae architecture in SSCs and progenitor spermatogonia. (**A–B**) Representative ultrastructure: Transmission electron micrographs showing representative mitochondrial ultrastructure in ID4-eGFP^Bright^ spermatogonial stem cells (SSCs; **A**) and ID4-eGFP^Dim^ progenitor spermatogonia (**B**). Scale bars, 200 nm. **(C–F) Mitochondrial size and shape:** Quantification of mitochondrial surface area (C; n = 77/79), length (D; n = 76/78), width (E; n = 77/80), and elongation expressed as length-to-width (L:W) ratio (F; n = 70/71). **(G) Cristae composition:** Relative distribution of lamellar, tubular and globular cristae in SSCs and progenitor spermatogonia (n = 337/471 cristae). **(H–J) Mitochondrial membrane complexity and dynamics:** Analysis of mitochondrial perimeter (H; n = 77/80), circularity (I; n = 67/69) and solidity (J; n = 66/67) as descriptors of mitochondrial membrane complexity and fusion–fission state. **(K–O) Cristae-specific architecture:** Quantification of cristae density, expressed as the ratio of total cristae area to mitochondrial surface area (K; n = 72/76 mitochondria), and of lamellar cristae width (L; n = 274/387), length (M; n = 276/410), elongation expressed as L:W ratio (N; n = 276/408) and junction width (O; n = 282/388 cristae). Values of n are given as SSCs/progenitors and refer to individual mitochondria (**C–F, H–K**) or cristae (**G, L–O**), following ROUT outlier removal (Q = 1%). Cells were pooled from n = 14 animals per group. Each data point represents an individual mitochondrion or lamellar crista, and horizontal lines denote the median. Statistical comparisons were performed using the Mann–Whitney U test. ***p < 0.001, ****p < 0.0001. Median and interquartile range values for all parameters are provided in **Supplementary Table S2**. **(P) Schematic ultrastructure:** Conceptual schematic summarising key ultrastructural features identified by quantitative TEM analysis. SSC mitochondria are characterised by wider lamellar cristae junctions and reduced lamellar cristae elongation, whereas progenitor mitochondria display increased cristae density and more elongated lamellar cristae, consistent with enhanced electron transport chain (ETC) complex assembly.

Analysis of cristae composition showed that lamellar cristae constituted the predominant cristae type in both SSCs and progenitors, with similar proportional distributions of lamellar, tubular, and globular cristae between groups (**Fig. 2G**). To assess mitochondrial membrane complexity and dynamic remodelling, additional shape descriptors were analysed (**Fig. 2H-J**). Mitochondrial perimeter, circularity, and solidity did not differ significantly between groups (*p* = 0.564; **Fig. 2H**), (*p* = 0.934; **Fig. 2I**), and (*p* = 0.162; **Fig. 2J**) respectively, suggesting no major differences in fusion-fission balance or membrane irregularity between SSCs and progenitor mitochondria.

TEM images revealed that progenitor mitochondria appeared consistently more electron-dense following osmium tetroxide staining, particularly within the inner membrane and cristae regions (**Fig. 2A-B**). Aligned with this, cristae-specific analyses revealed pronounced differences in inner membrane architecture (**Fig. 2K-O**). Progenitor mitochondria exhibited significantly higher cristae density, measured as the ratio of total cristae surface area to mitochondrial surface area (*p* < 0.0001; **Fig. 2K**). Lamellar cristae in progenitors were also wider and longer than those observed in SSCs, resulting in increased lamellar cristae elongation (*p* < 0.0001; **Fig. 2L**), (*p* < 0.0001; **Fig. 2M**), and (*p* = 0.0001; **Fig. 2N**), respectively. Conversely, lamellar cristae junction width was significantly reduced in progenitor mitochondria compared with SSCs (*p* < 0.0001; **Fig. 2O**).

In concert, these ultrastructural findings revealed cristae with wider lamellar junctions and reduced density in the stem cell population, while progenitors exhibited more elongated lamellar cristae at higher density (**Fig. 2P**). Narrower cristae junctions and increased cristae surface area in progenitors are consistent with enhanced capacity for ETC complex assembly and more efficient electron transport (Acharya et al., 2025; Cogliati et al., 2016).

### Proteomic profiling of mitochondrial membrane dynamics supports cristae organisation

To determine whether the differences in cristae organisation observed by TEM are reflected at the protein level, we examined the abundance of proteins regulating mitochondrial dynamics and cristae formation and maintenance (Skerrett-Byrne et al., 2026) (**Fig. 3**). Quantitative proteomics revealed selective differences in proteins associated with mitochondrial fission and inner membrane organisation between SSCs and progenitor spermatogonia.

**Figure 3.**
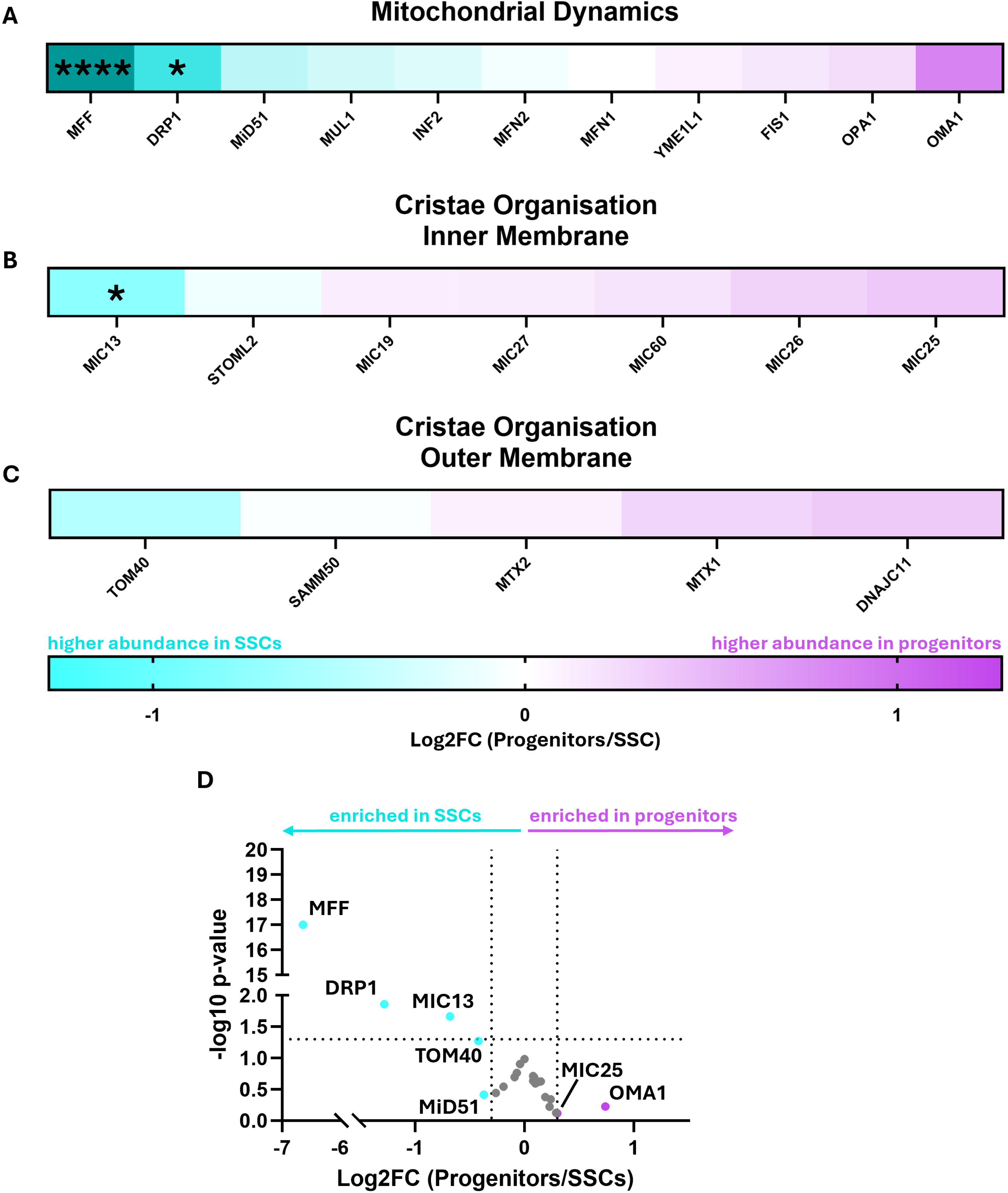
Proteomic analysis of proteins regulating mitochondrial dynamics and cristae organisation in SSCs and progenitor spermatogonia. (**A–C**) Heatmaps showing the log2 fold change (log2FC; Progenitors/SSCs) in abundance of proteins associated with mitochondrial dynamics (**A**), inner membrane cristae organisation (**B**), and outer membrane contact sites and cristae junction stability (**C**). Colour scale is symmetric around zero, ranging from cyan (higher abundance in SSCs) to purple (higher abundance in progenitors). **(D)** Volcano plot displaying the log2FC (Progenitors/SSCs) against the −log10 p-value for all quantified proteins within the mitochondrial dynamics and cristae organisation categories. Vertical dotted lines indicate the ≥|0.3| log2FC effect size threshold. The horizontal dotted line indicates p = 0.05. Proteins meeting the ≥|0.3| log2FC threshold are highlighted and labelled, with cyan indicating enrichment in SSCs and purple indicating enrichment in progenitors. All other proteins are shown in grey. All proteomic data are derived from n = 4 biological replicates. Statistical comparisons were performed using a paired two-tailed t-test. Statistical significance is indicated as *p < 0.05, ****p < 0.0001.

Among proteins regulating mitochondrial dynamics, the fission mediators DRP1 (–1.28 log_2_FC; *p* = 0.014) and mitochondrial fission factor (MFF; –6.64 log_2_FC; *p* < 0.0001) were significantly more abundant in SSCs (**Fig. 3A, D**). Notably, MFF was not detected in progenitor samples within our dataset. MFF functions as the primary outer membrane receptor for DRP1, facilitating its recruitment to fission sites (Koirala et al., 2013; Losón et al., 2013). Their coordinated enrichment in SSCs points to a greater capacity for mitochondrial fission and membrane remodelling in the stem cell population. In contrast, proteins associated with mitochondrial fusion, namely MFN1, MFN2, OPA1, and YME1L1, did not meet the ≥|0.3| log2FC threshold, and none reached statistical significance. OMA1 (0.74 log_2_FC), a stress-activated protease that cleaves OPA1 to drive inner membrane remodelling under mitochondrial stress (Gilkerson et al., 2021), showed enrichment in progenitors exceeding the effect size threshold but did not reach statistical significance (*p* = 0.22). Together, the absence of strong differences in fusion-related proteins is consistent with the lack of substantial differences in gross mitochondrial morphology observed by TEM (**Fig. 2C-F**).

Proteomic analysis of inner mitochondrial membrane structural components further revealed differential abundance of proteins involved in cristae organisation. MIC13 (–0.68 log_2_FC, *p* = 0.022) was significantly enriched in SSCs. MIC13 is a core MICOS subunit with a specific role in cristae junction formation, distinct from the broader scaffolding functions of MIC25, MIC26, MIC27, and MIC60 (Guarani et al., 2015), none of which met the ≥|0.3| log2FC threshold (**Fig. 3B, D**). The enrichment of MIC13 in SSCs, in the context of the wider cristae junctions observed by TEM, suggests that differences in cristae junction composition may contribute to the distinct inner membrane architecture that distinguishes SSCs from progenitor spermatogonia.

Among proteins associated with outer membrane contact sites, TOM40 (–0.42 log_2_FC, *p* = 0.054) showed a trend toward increased abundance in SSCs, while DNAJC11 (0.29 log2FC), MTX1, MTX2, and SAMM50 did not meet the ≥|0.3| log2FC threshold (**Fig. 3C**). The absence of significant differences in outer membrane-associated proteins suggests that the ultrastructural differences observed between SSCs and progenitors reflect remodelling of the inner membrane specifically, rather than broad changes in outer membrane architecture.

Collectively, these proteomic findings are consistent with the ultrastructural data and indicate that SSC mitochondria are characterised by enrichment of fission mediators and a distinct cristae junction composition, pointing to differences in both membrane remodelling capacity and inner membrane organisation between SSCs and progenitor spermatogonia.

### SSCs display enhanced mitochondrial respiratory capacity and metabolic plasticity relative to progenitor spermatogonia

Given the central role of cristae architecture in determining mitochondrial respiratory capacity, we next assessed whether these structural differences were associated with functional alterations in cellular energy metabolism in stem versus progenitor cells of the germline. Cellular bioenergetics were analysed in FACS-isolated SSCs and progenitor spermatogonia using Seahorse extracellular flux assays (**Fig. 4**). These assays were performed under normoxic culture conditions (21% O_2_), following an overnight *in vitro* incubation (at 10% O_2_), which differ considerably from the hypoxic SSC niche *in vivo* (<1.3% O_2_) (Bernstein et al., 2023; Morimoto et al., 2021). This distinction is important for interpreting the data, as the metabolic profiles reported here reflect the behaviour of these populations when oxygen availability is not limiting.

**Figure 4.**
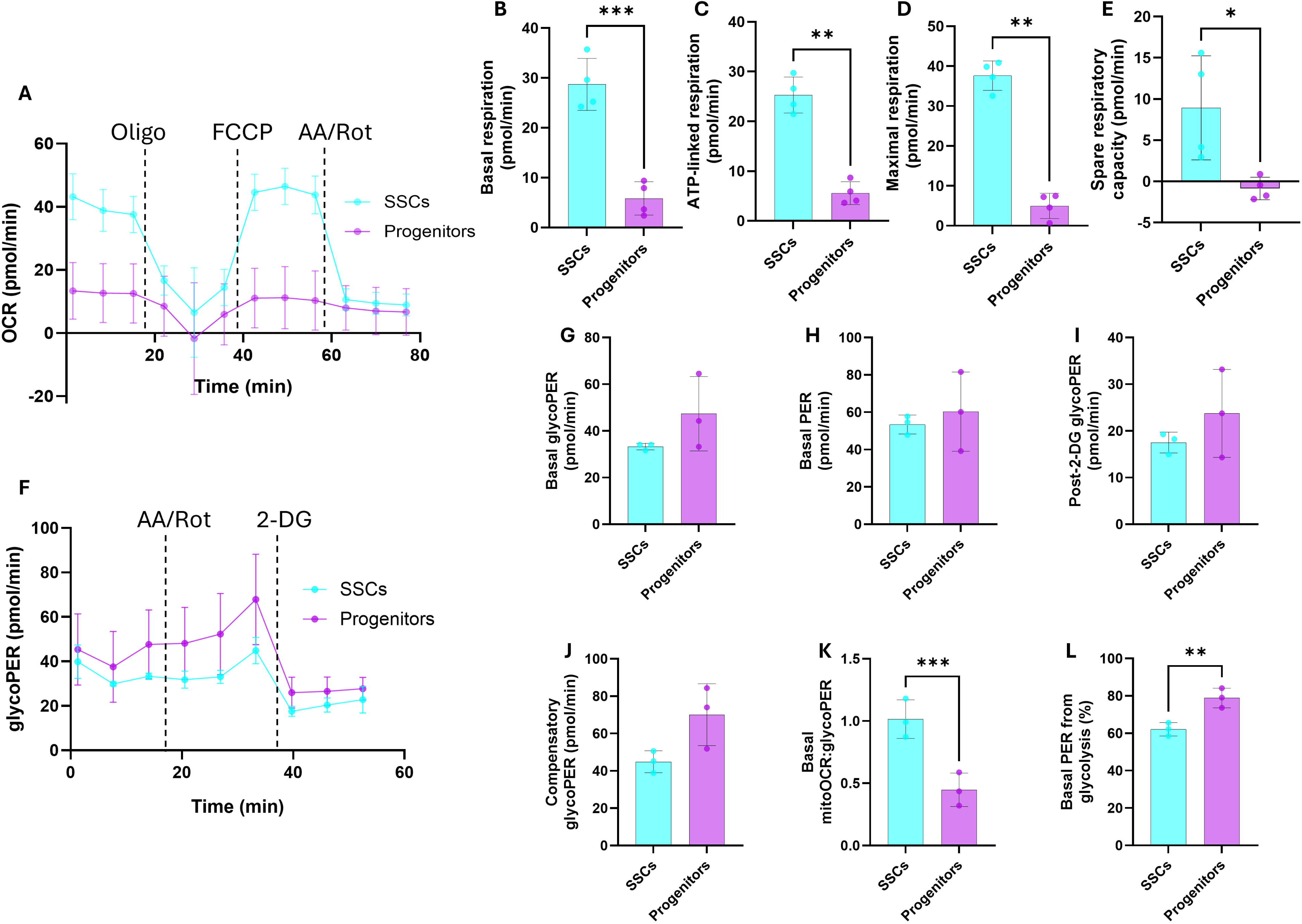
Mitochondrial respiratory capacity and glycolytic flux in SSCs and progenitor spermatogonia. (**A**) Real-time trace of oxygen consumption rate (OCR) in SSCs and progenitor spermatogonia during a Mito Stress Test (n = 4 biological replicates). Vertical lines indicate sequential injection of oligomycin (Oligo; ATP synthase inhibitor), FCCP (mitochondrial uncoupler), and antimycin A/rotenone (AA/Rot; complex III and I inhibitors), respectively. **(B–E)** Quantification of mitochondrial respiratory parameters derived from the Mito Stress Test, including basal respiration (**B**), ATP-linked respiration (**C**), maximal respiration (**D**), and spare respiratory capacity (**E**). **(F)** Real-time trace of the glycolytic proton efflux rate (glycoPER) during a Glycolytic Rate Assay (n = 3 biological replicates). Vertical lines indicate sequential injection of antimycin A/rotenone (AA/Rot) and 2-deoxy-D-glucose (2-DG; glycolytic inhibitor), respectively. **(G–J)** Quantification of glycolytic parameters derived from the Glycolytic Rate Assay, including basal glycolysis (**G**), basal total proton efflux rate (PER; **H**), post-2-DG acidification (**I**), and compensatory glycolysis (**J**). Basal glycolysis (**G**), compensatory glycolysis (**J**) and post-2-DG acidification (**I**) correspond to the final basal, maximal post-AA/Rot, and post-2-DG timepoints of the trace shown in (**F**), respectively. **(K–L)** Relative metabolic pathway reliance, expressed as the ratio of mitochondrial OCR to glycoPER (mitoOCR:glycoPER; **K**) and the percentage of total PER derived from glycolysis (**L**). All assays were performed under normoxic conditions following overnight culture. Data are presented as mean ± SD. Each data point represents an independent biological replicate. Statistical comparisons were performed using a paired two-tailed t-test. Statistical significance is indicated as *p < 0.05, **p < 0.01, and ***p < 0.001. Summary statistics for all parameters are provided in **Supplementary Table S3**.

#### Mitochondrial respiration is substantially elevated in SSCs under normoxic conditions

Real-time oxygen consumption rate (OCR) measurements are shown in **Fig. 4A–E**. ATP-linked respiration, calculated as the oligomycin-sensitive fraction of basal OCR, was significantly elevated in SSCs compared with progenitors (*p* = 0.0053; **Fig. 4C**), indicating a greater mitochondrial contribution to ATP production within the stem cell population. Basal respiration was also significantly higher in SSCs (*p* = 0.0003; **Fig. 4B**), though this metric reflects total cellular oxygen consumption and is therefore less specific to mitochondrial ATP synthesis than the oligomycin-sensitive fraction.

The most striking difference between populations was observed following FCCP-mediated uncoupling. Maximal respiration was markedly higher in SSCs compared with progenitors (*p* = 0.0012; **Fig. 4D**), demonstrating an enhanced capacity of SSC mitochondria to increase respiratory activity under conditions of elevated energetic demand. Consequently, spare respiratory capacity, defined as the difference between maximal and basal respiration, was significantly greater in SSCs (*p* = 0.0438; **Fig. 4E**), while progenitors displayed minimal reserve, indicating that progenitor mitochondria are operating close to their respiratory ceiling under these conditions. Together, the elevated maximal respiration and spare respiratory capacity of SSCs reveal a population with considerable mitochondrial reserve, a hallmark of metabolic plasticity.

#### Glycolytic flux is comparable between SSCs and progenitor spermatogonia

To examine glycolytic metabolism, proton efflux rate (PER) was measured using the Glycolytic Rate Assay (**Fig. 4F**). Under basal assay (normoxic) conditions, total glycolytic flux (glycoPER; *p* = 0.24; **Fig. 4G**), total PER (*p* = 0.58; **Fig. 4H**), and post-2-deoxy-D-glucose (2-DG) acidification (*p* = 0.28; **Fig. 4I**) did not differ significantly between SSCs and progenitors. Compensatory glycolysis following mitochondrial inhibition also showed no statistically significant difference between populations (*p* = 0.08; **Fig. 4J**), indicating broadly comparable glycolytic capacity in both cell states.

Despite comparable absolute glycolytic flux, the relative contribution of each metabolic pathway differed markedly between populations. The mitoOCR:glycoPER ratio was significantly higher in SSCs (*p* = 0.0005; **Fig. 4K**), indicating a greater relative reliance on mitochondrial respiration in normoxia. Conversely, the percentage of PER derived from glycolysis was significantly higher in progenitors (*p* = 0.003; **Fig. 4L**), indicating that extracellular acidification in progenitors was driven more strongly by glycolytic metabolism.

Collectively, these data demonstrate that when oxygen availability is not restricted, SSCs exhibit remarkable metabolic plasticity, maintaining glycolytic flux comparable to progenitors while simultaneously displaying substantially enhanced mitochondrial respiratory capacity and reserve.

#### Restoring hypoxia-inducible factor signalling suppresses mitochondrial respiration in cultured spermatogonia

Given that glycolytic metabolism has been shown to drive self-renewal in SSCs in multiple independent studies (Bernstein et al., 2023; Helsel et al., 2017a; Kanatsu-Shinohara et al., 2016; Morimoto et al., 2021), the extensive oxidative capacity of SSCs observed here was somewhat unexpected. We therefore hypothesised that this capacity is latent, and is normally restrained by hypoxia-responsive signalling in the niche. Because methodological constraints required our bioenergetic measurements to be made at atmospheric oxygen, we restored hypoxia signalling in primary cultures of undifferentiated spermatogonia using the prolyl hydroxylase inhibitor Daprodustat. We have previously shown that Daprodustat treatment restores expression of the hypoxia-inducible factor EPAS1, which is otherwise degraded in spermatogonia *in vitro*, and significantly increases the number of SSC-derived colonies following transplantation (Bernstein et al., 2023). After 24 h of treatment, Daprodustat reduced basal mitochondrial OCR by approximately 70% relative to vehicle-treated controls (39.8 ± 7.9 vs 11.7 ± 6.7 pmol/min; p = 0.0012; **Supplementary Fig. S4A, C**), whereas basal glycoPER was unchanged (90.5 ± 35.2 vs 106.1 ± 73.4 pmol/min; p = 0.54; **Supplementary Fig. S4B, D**). Accordingly, the proportion of PER derived from glycolysis increased (76.7 ± 9.1% vs 93.2 ± 3.0%; p = 0.0095; **Supplementary Fig. S4F**), and the mitoOCR:glycoPER ratio decreased approximately four-fold (0.53 ± 0.27 vs 0.13 ± 0.06; p = 0.018; **Supplementary Fig. S4G**). Compensatory glycolysis after mitochondrial inhibition tended to be lower with Daprodustat treatment, although the difference was not statistically significant (238.8 ± 121.0 vs 147.0 ± 93.9 pmol/min; p = 0.056; **Supplementary Fig. S4E**).

Together, these data indicate that restoring HIF signalling under normoxic conditions suppresses mitochondrial respiration and shifts spermatogonia toward glycolytic energy production. This supports the interpretation that the reserve oxidative capacity of SSCs is restrained by the hypoxic niche, and that this metabolic plasticity may be a defining and functionally important feature of the stem cell state.

### Proteomic profiling reveals pathway-specific differences in glycolytic enzymes between SSCs and progenitor spermatogonia

To determine whether the metabolic signatures observed by Seahorse analysis were reflected at the proteomic level, we examined the abundance of enzymes involved in glycolysis and associated metabolic pathways (**Fig. 5**). Relative changes in protein abundance (log_2_FC) across the glycolytic pathway are summarised in **Fig. 5A** with pathway mapping shown in **Fig. 5B**.

**Figure 5.**
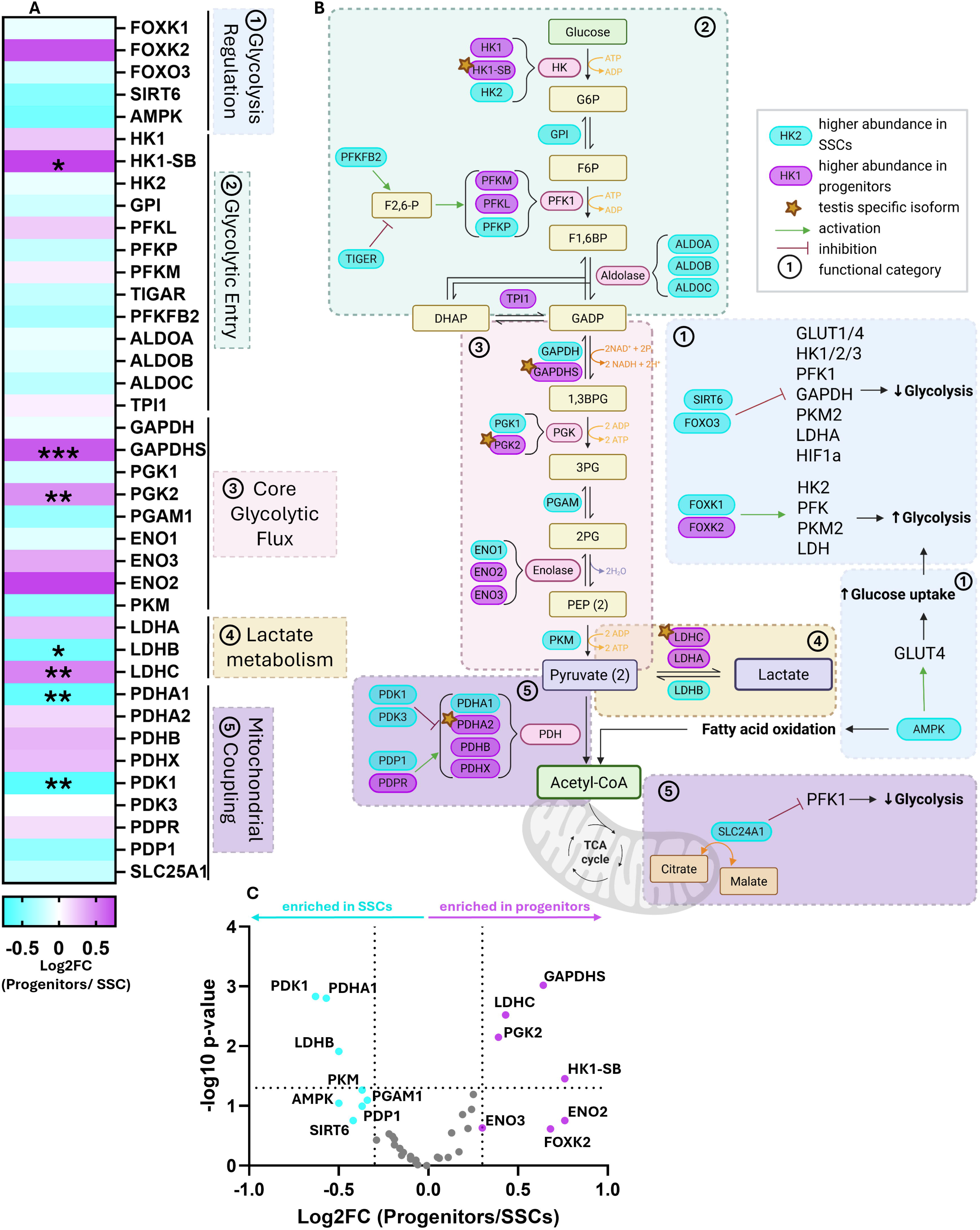
Proteomic profiling reveals pathway-specific differences in glycolytic enzyme abundance between SSCs and progenitor spermatogonia. (**A**) Heatmap showing log2FC (Progenitors/SSCs) of glycolysis and pyruvate metabolism– associated proteins identified by quantitative proteomics. Proteins are grouped according to functional categories: (1) glycolytic regulation, (2) glycolytic entry, (3) core glycolytic flux, (4) lactate metabolism, and (5) mitochondrial coupling. Colour scale is symmetric around zero, with cyan indicating higher abundance in SSCs and purple indicating higher abundance in progenitors. **(B)** Schematic representation of the glycolytic pathway with proteins detected in this dataset overlaid to illustrate relative enrichment and potential shifts in glycolytic flux and pyruvate fate. Numbered coloured boxes correspond to the functional categories in panel A. Refer to the in-figure key for symbol and colour definitions. **(C)** Volcano plot displaying log2FC (Progenitors/SSCs) against −log10 p-value for all quantified glycolysis– and pyruvate metabolism–associated proteins. Vertical dotted lines indicate the ≥|0.3| log2FC effect size threshold and the horizontal dotted line indicates p = 0.05. Proteins meeting the ≥|0.3| log2FC effect size threshold are highlighted in cyan (enriched in SSCs) or purple (enriched in progenitors) and labelled. All other proteins are shown in grey. All proteomic data are derived from n = 4 biological replicates. Statistical comparisons were performed using a paired two-tailed t-test. Statistical significance is indicated as *p < 0.05, **p < 0.01, ***p < 0.001.

Several enzymes associated with glycolytic flux were more abundant in progenitor spermatogonia. These included testis-specific glycolytic isoforms such as GAPDHS (0.64 log_2_FC, *p* < 0.001), PGK2 (0.39 log_2_FC, *p* = 0.007), and LDHC (0.43 log_2_FC, *p* = 0.003) (**Fig. 5C**). The increased abundance of these proteins is consistent with their established roles in later stages of germ cell differentiation and sperm function (Liu et al., 2019). The testis-specific hexokinase isoform HK1-SB (0.76 log_2_FC, *p* = 0.035) was also significantly higher in progenitors, indicating differences in the early steps of glycolytic entry between the two populations. The enolase isoforms ENO2 (0.76 log_2_FC) and ENO3 (0.3 log_2_FC), which catalyse the penultimate step in glycolysis, were similarly enriched in progenitors (**Fig. 5B**). In addition, the transcription factor FOXK2 (0.68 log_2_FC), a positive regulator of aerobic glycolysis through the upregulation of core glycolytic genes (Sukonina et al., 2019), showed higher abundance in progenitors.

In contrast, several enzymes associated with mitochondrial pyruvate metabolism were more abundant in SSCs. The pyruvate dehydrogenase (PDH) complex subunit PDHA1 (–0.57 log_2_FC, *p* = 0.0016) and its regulatory kinase PDK1 (–0.63 log_2_FC, *p* = 0.0014) were both significantly enriched in SSCs (**Fig. 5C**). Notably, LDHB (–0.5 log_2_FC, *p* = 0.012), which preferentially catalyses the conversion of lactate back to pyruvate (Vesell, 1965), was also enriched in SSCs, a finding that points to bidirectional pyruvate-lactate interconversion in SSCs and adds further support to their metabolic plasticity. Several additional metabolic regulators showed modest enrichment in SSCs above the ≥|0.3| log2FC threshold, including PKM (–0.37 log_2_FC), AMPK (–0.5 log_2_FC), and SIRT6 (–0.42 log_2_FC).

Mapping these proteins onto the glycolytic pathway revealed that enzymes enriched in progenitors predominantly participate in core glycolytic reactions generating ATP and metabolic intermediates, whereas proteins enriched in SSCs were more frequently associated with pyruvate entry into mitochondrial metabolism (**Fig. 5B**). Together, these patterns indicate a greater representation of proteins supporting mitochondrial pyruvate utilisation in SSCs relative to progenitors, consistent with the distinct metabolic phenotypes observed in extracellular flux assays. To further support these proteomic observations, QSM modelling of the dataset predicted greater lactate uptake capacity in SSCs relative to progenitors (**Supplementary Fig. S5**), consistent with the enrichment of LDHB and the capacity for bidirectional pyruvate-lactate interconversion in the stem cell population.

### Proteomic analysis reveals coordinated differences in oxidative phosphorylation components between SSCs and progenitor spermatogonia

Given the increased mitochondrial respiratory activity observed in SSCs under normoxic conditions, we next examined the abundance of proteins associated with OXPHOS and the mitochondrial electron transport chain (ETC) (**Fig. 6**). Proteomic mapping of ETC components across respiratory complexes I–V revealed selective differences in protein abundance between SSCs and progenitor spermatogonia (**Fig. 6A–E**). Within Complex I, NDUFB4 (0.39 log_2_FC, *p* = 0.04) was enriched in progenitors, while NDUFB1 (–0.75 log_2_FC, *p* = 0.01) showed higher abundance in SSCs, with remaining subunits showing only minor differences between populations.

**Figure 6.**
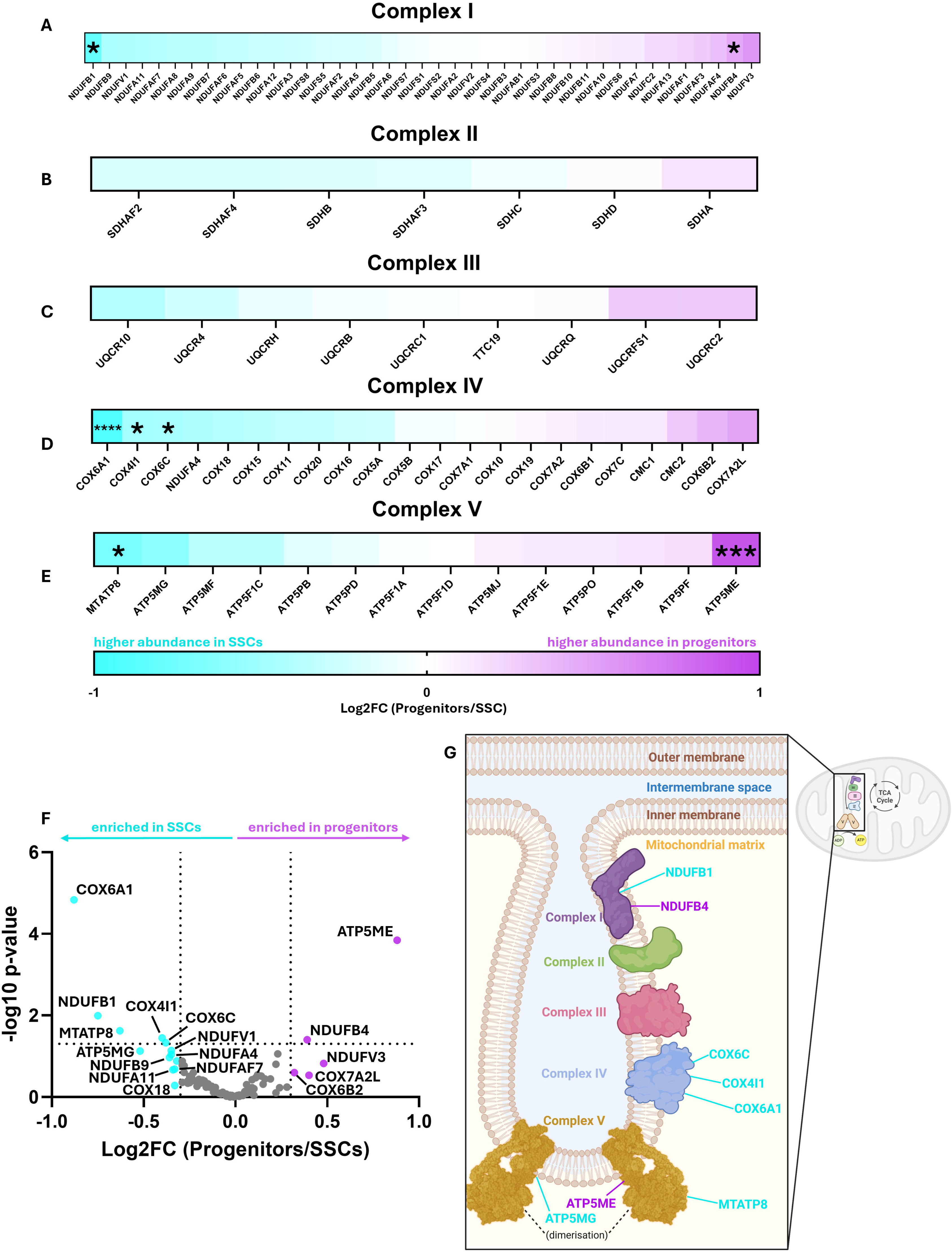
Proteomic analysis of electron transport chain components in SSCs and progenitor spermatogonia. (**A–E**) Heatmaps showing log2FC (Progenitors/SSCs) in abundance of proteins associated with mitochondrial respiratory Complex I (**A**), Complex II (**B**), Complex III (**C**), Complex IV (**D**), and Complex V (**E**). Colour scale is symmetric around zero, with cyan indicating higher abundance in SSCs and purple indicating higher abundance in progenitors. **(F)** Volcano plot displaying log2FC (Progenitors/SSCs) against −log10 p-value for all quantified ETC-associated proteins. Vertical dotted lines indicate the ≥|0.3| log2FC effect size threshold and the horizontal dotted line indicates p = 0.05. Proteins meeting the ≥|0.3| log2FC threshold are highlighted and labelled, with cyan indicating enrichment in SSCs and purple indicating enrichment in progenitors. All other proteins are shown in grey. **(G)** Schematic representation of the inner mitochondrial membrane illustrating the localisation of significantly enriched ETC subunits identified in this dataset. Proteins enriched in SSCs (cyan) and progenitors (purple) are indicated at their respective positions within Complexes I– V. All proteomic data are derived from n = 4 biological replicates. Statistical comparisons were performed using a paired two-tailed t-test. Statistical significance is indicated as *p < 0.05, ***p < 0.001, and ****p < 0.0001.

Among Complex IV subunits, COX6A1 (–0.88 log_2_FC, *p* < 0.0001), COX6C (–0.38 log_2_FC, *p* = 0.047), and COX4I1 (–0.4 log_2_FC, *p* = 0.036) were all significantly enriched in SSCs (**Fig. 6D, F**). These subunits play key roles in the terminal step of electron transfer to oxygen, facilitating efficient proton pumping and ATP production via OXPHOS (Čunátová et al., 2020; Pham et al., 2024).

Within Complex V, the ATP synthase subunit ATP5ME (0.88 log_2_FC, *p* = 0.0001) was significantly enriched in progenitors, while MTATP8 (–0.63 log_2_FC; *p* = 0.024) was significantly enriched in SSCs, and ATP5MG (–0.52 log_2_FC; *p* = 0.07) showed enrichment in SSCs above the ≥|0.3| log2FC threshold (**Fig. 6E, F**). Notably, ATP5ME and ATP5MG have been implicated in the formation and stabilisation of ATP synthase dimers that localise to regions of high membrane curvature at the apex of mitochondrial cristae (Hahn et al., 2016; Strauss et al., 2008) (**Fig. 6G**). In this context, differential abundance of ATP synthase-associated proteins may relate to the ultrastructural differences in cristae architecture observed by TEM, where progenitor spermatogonia displayed longer, wider lamellar cristae at higher density compared with SSCs (**Fig. 2K–M**).

Collectively, these data indicate that SSCs and progenitor spermatogonia differ selectively in the abundance of specific ETC subunits rather than showing global shifts across entire complexes. Together with the glycolytic proteomic data, this supports a model in which SSCs exist in a comparatively low energy-demand state in the niche, while retaining the plasticity to increase ATP production when stimulated, consistent with the functional metabolic analyses discussed above.

### Proteomic analysis of reactive oxygen species (ROS) production and redox balance

Given that increased mitochondrial respiration during spermatogonial differentiation has been associated with elevated reactive oxygen species (ROS) production (Chen et al., 2020b), we next examined the abundance of proteins involved in ROS generation and redox homeostasis between SSCs and progenitor spermatogonia (**Fig. 7**).

**Figure 7.**
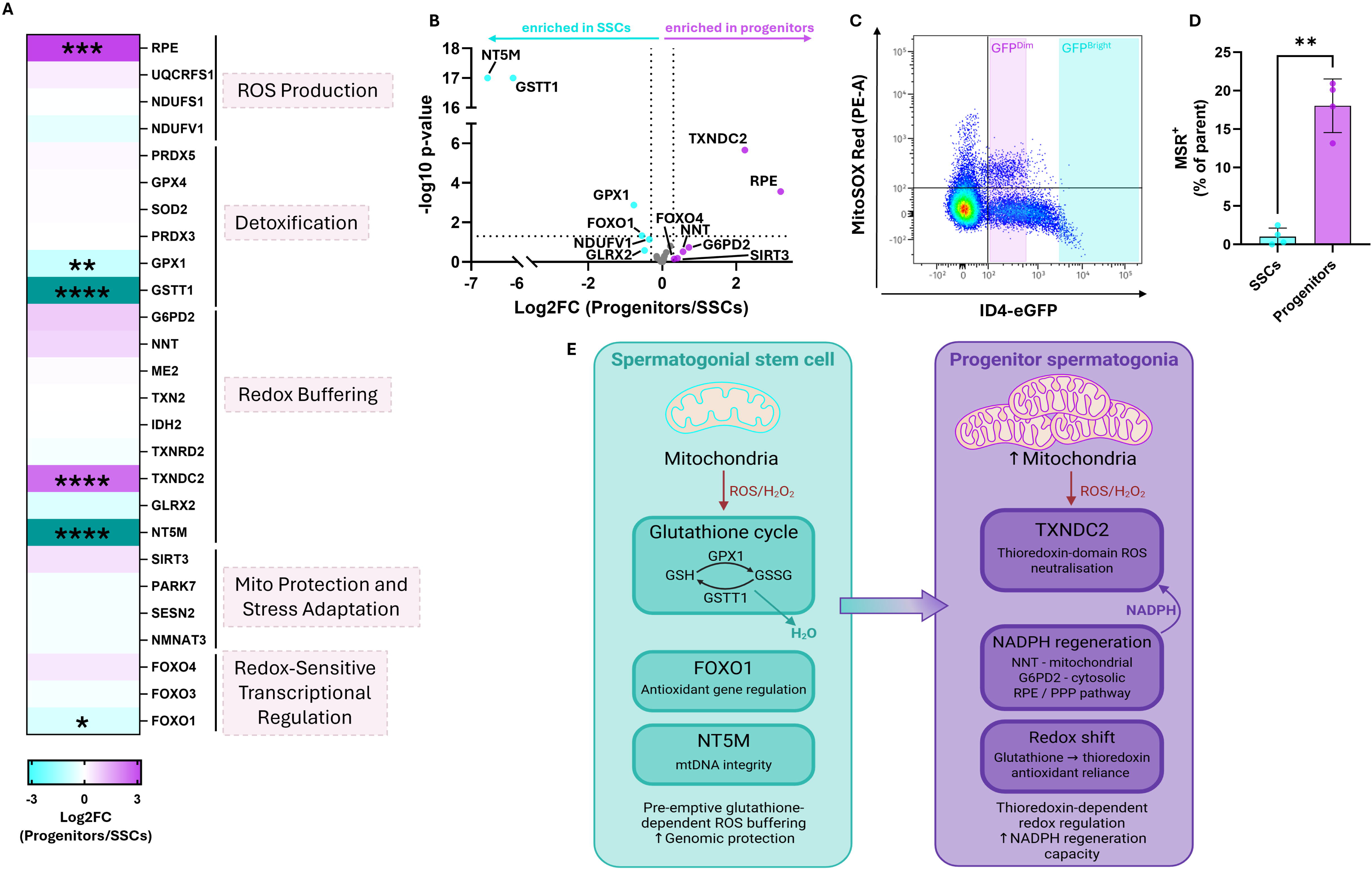
Proteomic and functional analysis of ROS production and redox balance in SSCs and progenitor spermatogonia. (**A**) Heatmap showing log2FC (Progenitors/SSCs) in abundance of proteins associated with ROS production, detoxification, redox buffering, mitochondrial protection and stress adaptation, and redox-sensitive transcriptional regulation. Colour scale is symmetric around zero (±3 log2FC), with cyan indicating higher abundance in SSCs and purple indicating higher abundance in progenitors. Proteins with abundance differences exceeding ±3 log2FC are represented by the darkest shade of the respective colour. **(B)** Volcano plot displaying log2FC (Progenitors/SSCs) against −log10 p-value for all quantified ROS– and redox-associated proteins. Vertical dotted lines indicate the ≥|0.3| log2FC effect size threshold and the horizontal dotted line indicates p = 0.05. Proteins meeting the ≥|0.3| log2FC threshold are highlighted and labelled, with cyan indicating enrichment in SSCs and purple indicating enrichment in progenitors. All other proteins are shown in grey. **(C)** Representative flow cytometry dot plot showing MitoSOX Red (MSR; PE-A) fluorescence against ID4-eGFP signal in live testicular cells (gating strategy in **Supplementary Fig. S1**). ID4-eGFP^Dim^ (progenitor-enriched) and ID4-eGFP^Bright^ (SSC-enriched) gates are highlighted in purple and cyan, respectively, and the horizontal line indicates the MSR⁺ gate. **(D)** Proportion of MSR⁺ events (% of parent) within ID4-eGFP^Bright^ (SSC) and ID4-eGFP^Dim^ (progenitor) populations. Each point represents one biological replicate. **(E)** Schematic summarising the distinct redox regulatory strategies identified in SSCs and progenitor spermatogonia. SSCs (cyan) rely predominantly on glutathione-dependent antioxidant machinery, including GPX1 and GSTT1, to neutralise mitochondrial ROS, supported by FOXO1-mediated antioxidant gene regulation and NT5M-dependent mitochondrial genome protection. Progenitor spermatogonia (purple) display a shift toward thioredoxin-dependent ROS neutralisation via TXNDC2, sustained by increased NADPH regeneration capacity through NNT, G6PD2, and the pentose phosphate pathway. The transition arrow indicates the shift in redox regulatory strategy accompanying the SSC-to-progenitor transition. Proteomic data (**A, B**) are derived from n = 4 biological replicates, and flow cytometry data (**C, D**) from n = 4 independent biological replicates (mean ± SD). All statistical comparisons were performed using paired two-tailed t-tests. *p < 0.05, **p < 0.01, ***p < 0.001, ****p < 0.0001.

Proteins associated with mitochondrial electron transport-derived ROS production were detected in both populations. Among these, NDUFV1 (–0.35 log_2_FC) showed modest enrichment above the ≥|0.3| log2FC threshold, while UQCRFS1 and NDUFS1 were detected in both populations below the threshold (**Fig. 7A**). In contrast, ribulose-phosphate 3-epimerase (RPE; 3.20 log_2_FC, *p* = 0.0003), a cytosolic enzyme of the pentose phosphate pathway (PPP), was significantly enriched in progenitors. As the PPP is a major source of cellular NADPH required to support antioxidant systems, this enrichment suggests differences in metabolic pathways that contribute to redox balance and suppression of oxidative stress (TeSlaa et al., 2023).

Proteins involved in ROS detoxification showed more pronounced variation between the two populations. Two enzymes of the glutathione pathway, GSTT1 (–6.08 log_2_FC, *p* < 0.0001) and GPX1 (–0.77 log_2_FC, *p* = 0.001) were significantly enriched in SSCs. GSTT1, which plays an important role in oxidative stress management, displayed one of the largest abundance changes among metabolic proteins identified in this dataset, while GPX1 catalyses the reduction of hydrogen peroxide using glutathione (Ito et al., 2011). Additional detoxification enzymes, including PRDX3, PRDX5, GPX4, and SOD2, were detected in both populations but did not meet the ≥|0.3| log2FC threshold.

Among proteins associated with cellular redox buffering and thiol-based antioxidant systems, NNT (0.56 log_2_FC) and G6PD2 (0.72 log2FC) showed enrichment in progenitors, while GLRX2 (–0.48 log2FC), a mitochondrial glutaredoxin involved in thiol-disulfide exchange and iron-sulphur cluster assembly (Lillig et al., 2005), showed enrichment in SSCs. ME2, TXN2, IDH2, and TXNRD2 were present in both populations below the threshold. In contrast, TXNDC2 (2.23 log_2_FC, *p* < 0.0001), a testis-enriched thioredoxin-domain protein implicated in germ cell redox regulation (Javadirad and Mokhtari, 2021), was significantly enriched in progenitors, suggesting potential differences in thioredoxin-dependent antioxidant systems between the two populations.

Proteins involved in mitochondrial stress adaptation and maintenance were also detected. Among these, the mitochondrial 5′-nucleotidase NT5M (–6.64 log_2_FC, *p* < 0.0001), which contributes to the regulation of mitochondrial nucleotide pools and mitochondrial genome stability (Rampazzo et al., 2000), was strongly enriched in SSCs and was not detected in the progenitor samples. The redox-responsive transcription factor FOXO1 (–0.55 log_2_FC, *p* = 0.047) was also significantly enriched in SSCs. SIRT3 (0.41 log2FC), a mitochondrial deacetylase involved in antioxidant defence and metabolic regulation via FOXO3a-dependent pathways (Sundaresan et al., 2009), showed enrichment in progenitors above the threshold, a finding that warrants further investigation given its established role in ROS buffering. PARK7, NMNAT3 and SESN2 were present in both populations but did not meet the ≥|0.3| log2FC threshold.

To determine whether the proteomic differences described above were accompanied by measurable differences in mitochondrial superoxide accumulation, we performed MitoSOX Red (MSR) flow cytometry (**Fig. 7C–D**). SSCs displayed a significantly lower proportion of MSR⁺ events than progenitors (1.0 ± 1.1% vs 18.0 ± 3.5%; *p* = 0.0045), indicating that SSCs maintain a lower basal mitochondrial superoxide state than their progenitor counterparts, consistent with the proteomic evidence for enhanced antioxidant and redox-buffering capacity in this population.

Together, these findings reveal selective differences in proteins associated with ROS detoxification, redox buffering, and mitochondrial maintenance between SSCs and progenitor spermatogonia, consistent with the lower basal mitochondrial superoxide burden observed in SSCs by MSR flow cytometry. In particular, the strong enrichment of glutathione-dependent antioxidant enzymes and mitochondrial maintenance factors in SSCs, together with this functional evidence of reduced steady-state ROS, indicates a robust intrinsic antioxidant capacity in the stem cell population, consistent with the need to preserve genomic integrity in self-renewing cells.

## DISCUSSION

Elucidating the functional regulation of stem cells has important implications for the development of therapeutic and conservation strategies that manipulate fate decisions *in vitro*. In this study, we conducted a multimodal analysis of mitochondrial structure, metabolic activity and redox regulation in mouse SSCs and progenitor spermatogonia to determine whether transition to a progenitor state is accompanied by metabolic remodelling. Previous studies investigating similar parameters have employed retinoic acid-induced differentiation *in vitro* or isolation strategies that do not allow clear separation of progenitor and differentiating spermatogonia (Chen et al., 2020b; Zhang et al., 2024). By combining highly purified populations (Chan et al., 2014; Helsel et al., 2017b) with complementary structural, functional and proteomic approaches, we show that the transition from SSCs to progenitor spermatogonia is associated with coordinated changes in mitochondrial abundance, cristae organisation, metabolic activity and redox regulation. These findings suggest that the transition from the stem cell to the progenitor state involves mitochondrial and metabolic remodelling that precedes the larger bioenergetic shifts observed during later stages of germ cell development (Varuzhanyan and Chan, 2020). Contrastingly, SSCs demonstrate a remarkable capacity for metabolic plasticity, likely reflecting their requirement for flexibility in response to environmental cues such as steady state versus regenerative conditions (**Fig. 8**).

**Figure 8.**
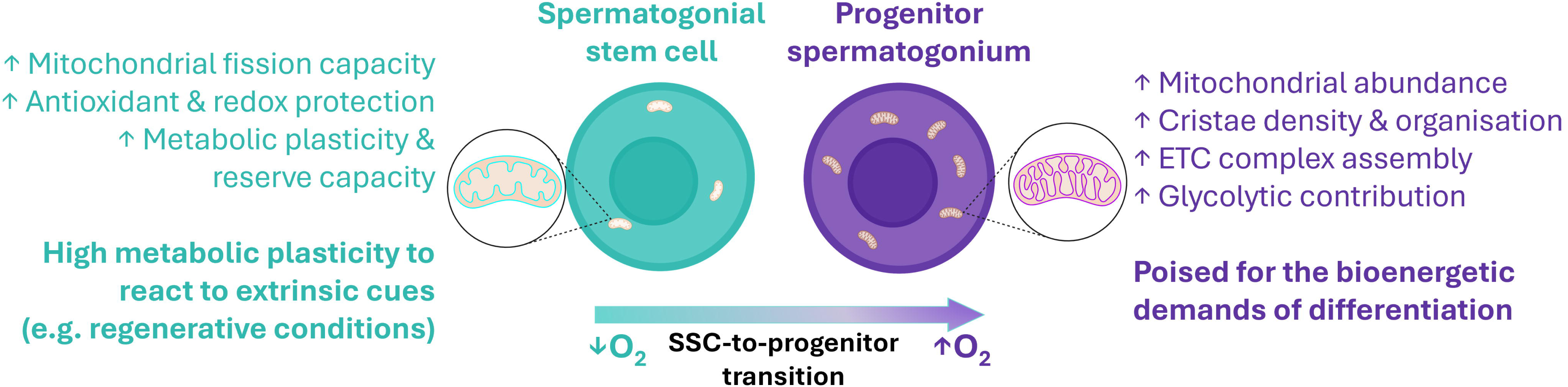
Schematic summary of the mitochondrial and metabolic features that distinguish SSCs from progenitor spermatogonia. Spermatogonial stem cells (SSCs; cyan) reside in a hypoxic niche and are characterised by enhanced mitochondrial fission capacity, robust antioxidant and redox protection, and high metabolic plasticity reflected by substantial respiratory reserve. Mitochondrial insets illustrate the sparse cristae density and wider cristae junctions characteristic of SSC mitochondria. Progenitor spermatogonia (purple) display increased mitochondrial abundance, higher cristae density and organisation, greater ETC complex assembly capacity, and a higher relative glycolytic contribution. Mitochondrial insets illustrate the higher cristae density and more elongated lamellar cristae characteristic of progenitor mitochondria. The gradient arrow indicates the SSC-to-progenitor transition, accompanied by increasing oxygen availability as cells move along the basement membrane of the seminiferous tubule. Schematic is conceptual and not drawn to scale.

Consistent with observations in other stem cell systems, the SSC-to-progenitor transition was associated with increased mitochondrial abundance, reflected by both increased mitochondrial DNA copy number and higher mitochondrial fluorescence intensity (**Fig. 1A, B**). Expansion of the mitochondrial network is a common feature of stem cell differentiation and often reflects a gradual shift toward greater mitochondrial contribution to cellular energy metabolism (Bahat and Gross, 2019; Wanet et al., 2015). Similar increases in mitochondrial copy number have been reported during differentiation of embryonic and pluripotent stem cells, where mitochondrial biogenesis accompanies the transition from a quiescent stem cell state to a more metabolically active progenitor phenotype (Cho et al., 2006; Facucho-Oliveira et al., 2007).

Despite this increase in mitochondrial abundance, TEM analyses indicate that mitochondrial remodelling during the SSC-to-progenitor transition occurs primarily at the level of cristae organisation rather than through changes in gross mitochondrial morphology. Progenitor mitochondria exhibited significantly higher cristae density together with wider and longer lamellar cristae, indicating increased inner membrane complexity, whereas SSC mitochondria displayed reduced cristae packing and less elongated lamellar cristae (**Fig. 2K–N**). Importantly, these differences occurred in the absence of major changes in mitochondrial size, elongation, circularity, or solidity (**Fig. 2C–F, H–J**), suggesting that overall mitochondrial shape and fusion-fission capacity remain broadly similar between the two populations. Consistent with these structural observations, the visibly darker appearance of progenitor mitochondria following osmium tetroxide staining likely reflects increased lipid membrane content within the organelle, given osmium’s affinity for lipid membranes (Belazi et al., 2009), and is in line with the higher cristae density observed by TEM. Similar features have been described in other stem cell systems, where immature stem cell mitochondria are typically small and characterised by poorly developed cristae and an electron-lucent matrix, while differentiated cells exhibit elongated mitochondria with abundant cristae and a more electron-dense matrix (Cho et al., 2006; Varum et al., 2011).

Cristae organisation is a key determinant of mitochondrial bioenergetic efficiency. Tightly folded lamellar cristae increase the surface area available for ETC complexes and ATP synthase, thereby enhancing ATP production (Afzal et al., 2021). At the same time, excessive cristae packing may introduce diffusional constraints for metabolites such as ADP within the cristae lumen (Cogliati et al., 2016; Mannella, 2006). The comparatively reduced cristae density observed in SSCs may therefore represent a structural configuration compatible with lower basal respiratory activity while retaining the capacity for metabolic adaptation.

Variation in ATP synthase dimer angle has been associated with differences in cristae morphology and metabolic capacity across species, with wider dimer angles producing narrower lamellar cristae that may facilitate efficient oxygen utilisation in low-oxygen environments (Stachowiak et al., 2013). In this context, the narrower lamellar cristae observed in SSCs (**Fig. 2L**) may reflect structural adaptation to the hypoxic niche in which these cells reside, whereas the wider cristae in progenitors could be associated with spatial or developmental transitions toward regions of higher oxygen availability within the testis (Bernstein et al., 2023). Higher-resolution imaging will be required to directly resolve these structural parameters. Although direct quantification of cristae apex angle was not feasible at the resolution of the current dataset, proteomic data provide mechanistic insight consistent with these ultrastructural observations. In particular, ATP5ME was enriched in progenitors while ATP5MG and MTATP8 showed greater abundance in SSCs, with all three subunits implicated in ATP synthase dimer formation and cristae apex curvature (Hahn et al., 2016; Strauss et al., 2008). The opposing enrichment of these subunits between populations suggests that differences in ATP synthase dimer composition may contribute to the distinct cristae architectures observed (Buzzard et al., 2024), a relationship that higher-resolution approaches such as electron tomography will be required to directly resolve.

The enrichment of the MICOS component MIC13 in SSCs, together with wider cristae junctions observed by TEM, points to a distinct crista junction architecture in the stem cell population. Given emerging evidence that MICOS-dependent remodelling of crista junctions can influence mitochondrial membrane potential and metabolic responsiveness (Gottschalk et al., 2022; Katti et al., 2026; Tomar et al., 2023), differential MIC13 abundance may represent a mechanism through which SSCs modulate inner membrane organisation to support metabolic plasticity. In parallel, modest enrichment of OMA1 in progenitors suggests a greater capacity for stress-responsive inner membrane remodelling in this population (Gilkerson et al., 2021), consistent with their transition toward a more metabolically active state.

Classical mediators of mitochondrial fusion showed minimal differences between SSCs and progenitor spermatogonia, consistent with the absence of substantial changes in gross mitochondrial morphology observed by TEM. This contrasts with transcriptomic analyses of the same cell types reporting increased *Mfn1* expression in progenitors (Helsel et al., 2017b; Lord and Nixon, 2020), whereas our proteomic data indicate no change in MFN1 abundance between populations, and MFN2 did not meet the ≥|0.3| log2FC threshold (**Fig. 3A**). These discrepancies likely reflect post-transcriptional regulation of mitochondrial dynamics proteins, consistent with the poor transcript-protein correlation we have recently reported in spermatogonia (Skerrett-Byrne et al., 2026).

In contrast, SSCs exhibited increased abundance of mitochondrial fission regulators DRP1 and MFF (**Fig. 3A, D**), as well as the mitochondrial maintenance factor NT5M (**Fig. 7A-B**), suggesting an enhanced capacity for mitochondrial remodelling and quality control in the stem cell population (Basu et al., 2017; Nolden et al., 2023; Rampazzo et al., 2000). Such a profile is consistent with maintenance of a small, dynamic mitochondrial population, a feature commonly associated with stem cell states across multiple tissues (Bahat and Gross, 2019; Ito and Suda, 2014). The absence of detectable differences in mitochondrial elongation or morphology by TEM could reflect the dynamic and transient nature of fission events, which may not be fully captured by two-dimensional imaging (Acharya et al., 2026b; Kondadi et al., 2020).

A key finding of this study was the remarkable metabolic plasticity displayed by SSCs when removed from their hypoxic niche and cultured under normoxic conditions. Rather than representing a limitation of the experimental approach, this methodological feature revealed a latent oxidative capacity in SSCs that is likely to be constrained under the hypoxic conditions of the niche, and that distinguishes them from progenitor spermatogonia. Consistent with this, restoring HIF signalling with Daprodustat suppressed mitochondrial respiration in cultured spermatogonia and shifted their energy production toward glycolysis (**Supplementary Fig. S4**). Under normoxia, SSCs exhibited substantially higher mitochondrial respiration, greater ATP-linked respiratory output, and markedly increased spare respiratory capacity relative to progenitors, while maintaining comparable absolute glycolytic flux (**Fig. 4**). This profile is consistent with a population that operates under metabolic restraint in its native hypoxic niche, where glycolytic metabolism supports self-renewal (Helsel et al., 2017a; Kanatsu-Shinohara et al., 2016), yet retains considerable functional respiratory reserve that can be rapidly mobilised in response to changes in oxygen availability or energetic demand. This capacity may facilitate swift transitions between quiescence and activation as circumstances demand (Tang and Rando, 2014).

Such metabolic plasticity is increasingly recognised as a defining feature of adult stem cell populations across tissues. In haematopoietic stem cells, the capacity to switch between glycolytic and oxidative metabolism underpins their ability to respond to regenerative demands while maintaining quiescence under homeostatic conditions (Chandel et al., 2016; Ito and Suda, 2014). Similarly, muscle satellite cells and neural stem cells retain oxidative reserve that is mobilised upon activation (Khacho et al., 2016; Tang and Rando, 2014). The present findings suggest that SSCs share this property, and that their metabolic plasticity may be particularly important in contexts requiring rapid germline regeneration, such as following genotoxic injury (Bernstein et al., 2023; La et al., 2022; Li et al., 2026). In progenitor spermatogonia, by contrast, the minimal spare respiratory capacity and greater relative glycolytic contribution suggest a population already operating closer to its bioenergetic ceiling, consistent with a transitional state poised for commitment to differentiation rather than niche re-entry.

Proteomic data provide further insight into the basis of these metabolic differences. The relative enrichment of glycolytic enzymes and transcriptional regulators of aerobic glycolysis, including FOXK2, in progenitors supports a shift towards greater glycolytic engagement during the SSC-to-progenitor transition. In contrast, the enrichment of proteins involved in mitochondrial pyruvate utilisation in SSCs, including PDHA1, PDK1 and LDHB, together with QSM predictions of greater lactate uptake capacity (**Supplementary Fig. S5D**), points to a population primed for bidirectional carbon substrate utilisation rather than commitment to a single metabolic pathway. Notably, these proteomic differences occur in the absence of widespread changes in ETC component abundance, indicating that the transition between these states does not involve global remodelling of OXPHOS machinery. However, selective enrichment of COX6A1, COX6C and COX4I1 in SSCs suggests differences in complex IV subunit composition that may influence electron transfer efficiency (**Fig. 6D, F**). Given the established sensitivity of Complex IV subunit composition to oxygen availability (Fukuda et al., 2007), this selective enrichment may reflect an adaptation of respiratory machinery to the hypoxic SSC niche, consistent with the metabolic plasticity observed functionally. These findings were perhaps unexpected in light of previous scRNA-seq studies reporting increased expression of glycolytic genes in SSCs and enrichment of OXPHOS pathways in progenitor and differentiating spermatogonia (Guo et al., 2017; Hermann et al., 2018; Lord and Nixon, 2020). However, it is important to appreciate that these studies did not delineate between progenitor and differentiating spermatogonia, limiting their ability to define the precise metabolic transition occurring at the SSC-to-progenitor boundary. Additionally, our recently published multiomic analysis has identified that correlation between transcript and protein abundance in spermatogonia is poor (Skerrett-Byrne et al., 2026), as has previously been reported in the testis as a whole (Jiang et al., 2020), highlighting that substantial post-transcriptional regulation of metabolic genes is likely. Together, these observations underscore the importance of proteomic approaches for accurately defining the molecular landscape of metabolic transitions in stem cell populations.

In addition to metabolic differences, proteomic profiling revealed distinct patterns of proteins associated with ROS detoxification and redox regulation between SSCs and progenitor spermatogonia. SSCs displayed strong enrichment of glutathione pathway enzymes, including GSTT1 and GPX1, and the mitochondrial glutaredoxin GLRX2, together with mitochondrial maintenance factors such as NT5M and the redox-responsive transcription factor FOXO1 (**Fig. 7A-B**). The functional importance of GSTT1 in the stem cell compartment is supported by our recent characterisation of the *Gstt1* knockout mouse, in which disruption of this gene impairs spermatogonial maintenance resulting in an increased number of Sertoli cell-only tubules (Skerrett-Byrne et al., 2026). Direct functional assessment of mitochondrial superoxide accumulation provided complementary support for this broader antioxidant signature. Notably, SSCs maintained a significantly lower proportion of MSR⁺ events than progenitors, indicating a lower steady-state mitochondrial superoxide burden in the stem cell population (**Fig. 7C–D**).

Considered together with the elevated spare respiratory capacity of SSCs described above, these findings suggest that SSCs do not simply respire less than progenitors, but actively decouple respiratory potential from oxidative output, holding substantial reserve capacity available while limiting steady-state ROS through pre-emptive glutathione-dependent buffering. This active redox strategy, rather than a passive response to oxidative stress, is consistent with the broader capacity for metabolic plasticity that characterises SSCs. Furthermore, this distinguishes SSCs from the canonical model of redox control in quiescent, niche-resident stem cells, in which low ROS is typically attributed to a metabolic preference for glycolysis over oxidative phosphorylation, thereby limiting oxidative ROS production at its source (Ushio-Fukai and Rehman, 2014). Instead, this reserve capacity appears to enable SSCs to upregulate mitochondrial respiration in response to changing oxygen availability while maintaining protection against the oxidative damage that would otherwise accompany increased respiratory activity. Such pre-emptive redox buffering aligns with the long-term self-renewal requirements of SSCs, where protection of genomic integrity is essential for sustained germline function (Jensen and Rothenberg, 2020), and may similarly support the robust antioxidant protection required for stem cell activation in regenerative contexts (La et al., 2022; Urish et al., 2009). This is consistent with a broader principle increasingly recognised across adult stem cell populations, in which low basal ROS functions as an active determinant of the self-renewing state (Bigarella et al., 2014; Ito and Suda, 2014).

By contrast, progenitor spermatogonia displayed enrichment of TXNDC2, a thioredoxin-domain protein implicated in germ cell redox regulation (Smith et al., 2013), and NNT, which regenerates NADPH to support mitochondrial antioxidant systems (Rydström, 2006), suggesting a shift in the balance between glutathione– and thioredoxin-dependent antioxidant systems and NADPH regeneration capacity as cells transition to the progenitor state. The enrichment of SIRT3 in progenitors further supports this shift, as SIRT3-mediated deacetylation of FOXO3a activates mitochondrial antioxidant gene expression independently of the glutathione pathway (Sundaresan et al., 2009). The opposing enrichment of the related transcription factors FOXO1 in SSCs and SIRT3 in progenitors, both redox-sensitive regulators operating through distinct FOXO family members, points to a coordinated remodelling of redox-sensitive transcriptional programmes as part of the broader cellular reprogramming underlying the SSC-to-progenitor transition, rather than acting solely as a protective response to metabolic change.

Taken together, our findings support a model in which SSCs maintain a metabolically restrained yet functionally competent mitochondrial network, characterised by lower mitochondrial abundance, reduced cristae density, and limited steady-state respiratory demand, while preserving considerable functional respiratory reserve, remodelling potential, and robust antioxidant protection. This combination of metabolic restraint and latent oxidative capacity is the hallmark of a population uniquely adapted to thrive in a hypoxic niche while retaining the plasticity to rapidly upregulate energy production in response to changing environmental demands. During the transition to progenitor spermatogonia, mitochondrial abundance increases and inner membrane architecture becomes more optimised for ATP production, as reflected by increased cristae density and elongation, alongside a shift toward greater metabolic engagement and glycolytic contribution. This coordinated remodelling of mitochondrial structure, metabolism, and redox regulation likely represents an early preparatory step that enables progenitor spermatogonia to meet the elevated energetic and biosynthetic demands of later stages of spermatogenesis. Overall, these findings establish metabolic plasticity as a defining feature of the spermatogonial stem cell state and provide new insight into the mitochondrial and metabolic adaptations that accompany the transition from stem cell to progenitor in the germline.

### Limitations of the study

The findings of this study should be considered in the context of several methodological considerations. Ultrastructural analyses were performed using two-dimensional TEM, which enables rigorous quantitative assessment of cristae density and morphology, however cannot fully capture the three-dimensional complexity of mitochondrial inner membrane organisation. Regardless, structural differences described here have been supported by complementary proteomic data, revealing meaningful differences between the two populations. Future studies using electron tomography or focused-ion-beam scanning electron microscopy will be valuable for more precisely defining cristae architecture and membrane curvature in SSCs and progenitor spermatogonia (Acharya et al., 2026b; Perkins et al., 1997).

A further consideration is that the extracellular flux analyses were performed under normoxic conditions following overnight culture, which differs from the hypoxic environment of the SSC niche *in vivo*. This deviation from physiological oxygen levels was instrumental in revealing the latent respiratory capacity of SSCs, although the extent to which this capacity is engaged *in vivo* remains to be determined. While our pharmacological HIF stabilisation using Daprodustat captures only part of the cellular response to low oxygen, these experiments support the interpretation that hypoxia-responsive signalling restrains this capacity. Assessing SSC bioenergetics across physiologically relevant oxygen tensions will be important to establish how this metabolic plasticity is deployed within the niche.

Finally, the spermatogonia used in this study were isolated from neonatal testes at postnatal days 6-8, due to their inherent abundance and purity at this developmental stage. Whether the metabolic plasticity demonstrated here is sustained in adult SSCs remains an important question, given emerging evidence that spermatogonial metabolism may differ between postnatal and adult stages (Voigt et al., 2022). Future experiments should therefore focus on determining whether the metabolic features identified here are conserved throughout postnatal development and into adulthood.

## Acknowledgements

Thank you to Nicole Cole and Dr Huiming Zhang from the University of Newcastle’s Central Analytical facilities for their assistance with FACS isolations and TEM imaging, respectively.

## Author contributions

Conceptualization, K.B.D. and T.L;

Methodology, K.B.D., J.J.F., S.A., and T.L.;

Formal Analysis K.B.D., J.J.F., S.A., and T.L.;

Investigation, K.B.D., J.J.F., S.A., and T.L.;

Resources, J.J.F., B.N., and T.L.;

Writing – Original Draft K.B.D. and T.L.;

Writing – Review & Editing K.B.D., J.J.F., S.A., B.N. and T.L.;

Supervision, J.J.F., B.N., and T.L.;

Funding Acquisition, B.N. and T.L.

## SUPPLEMENTAL FIGURE LEGENDS

**Supplementary Figure S1. Representative gating strategy for MitoSOX Red flow cytometry analysis.** Sequential gating strategy applied to testicular cell suspensions prior to identification of ID4-eGFP subpopulations. Cells were first gated on FSC-A/SSC-A to exclude debris and non-cellular events (**1**, “Cells”), followed by exclusion of non-viable cells based on SYTOX Deep Red fluorescence (**2**, “Live cells”; SYTOX⁻ gate indicated). Downstream identification of ID4-eGFPBright and ID4-eGFPDim subpopulations and MitoSOX Red-positive gating is shown in **Fig. 7C**.

**Supplementary Figure S2. Definition of lamellar cristae measurements derived from 2D TEM images.** Representative transmission electron micrograph of a spermatogonial mitochondrion illustrating lamellar cristae (left), with cristae manually segmented (blue) for quantitative analysis. The inset schematic (right) depicts the parameters used to quantify lamellar cristae morphology from 2D TEM sections, including maximum crista length, maximum crista width, and crista junction width at the point of connection to the inner mitochondrial membrane. Cristae density was calculated as the ratio of total cristae surface area to mitochondrial surface area. Measurements were performed only on clearly discernible lamellar cristae with well-defined membrane boundaries. Quantification was conducted in two dimensions and reflects the apparent cristae geometry within individual TEM sections. Scale bar = 200nm. Schematic is not to scale.

**Supplementary Figure S3.**Representative images of ID4-eGFP^Bright^ or ID4-eGFP^Dim^ cells seeded in XF96 cell culture microplates at 50,000 cells/well for Seahorse metabolic assays after an overnight incubation at 37°C in 10% O_2_ and 5% CO_2_. Scale bar = 300μm.

**Supplementary Figure S4. Daprodustat suppresses mitochondrial respiration and shifts cultured spermatogonia toward glycolysis.** Primary cultures of undifferentiated spermatogonia were treated with 125 µM Daprodustat (DStat) or DMSO vehicle for 24 h at 10% O_2_, before the Seahorse Glycolytic Rate Assay at 21% O_2_. (**A–B**) Kinetic traces of oxygen consumption rate (OCR; **A**) and glycolytic proton efflux rate (glycoPER; **B**), shown as mean ± SD. Dashed lines indicate the injections of antimycin A/rotenone (AA/Rot) and 2-deoxy-D-glucose (2-DG). (**C–G**) Basal mitochondrial OCR (mitoOCR; C), basal glycoPER (**D**), compensatory glycoPER (**E**), percentage of basal PER derived from glycolysis (**F**), and ratio of basal mitoOCR to glycoPER (**G**). Bars show mean ± SD, and each point is one biological replicate (an independent culture; mean of 2–3 technical wells). Statistical comparisons were performed using paired two-tailed t-tests (n = 5). *p < 0.05, **p < 0.01.

**Supplementary Figure S5. Quantitative systems metabolism (QSM) modelling of central carbon metabolism in SSCs and progenitor spermatogonia.** (A) Simplified schematic summarising predicted metabolic trends inferred from proteomic data. Progenitor spermatogonia show relatively greater representation of glycolytic enzymes, whereas SSCs display a tendency toward increased lactate uptake from the seminiferous microenvironment. (**B–D**) QSM modelling outputs showing predicted maximal ATP production capacity and substrate uptake capacities derived from protein abundance values. (**B**) Predicted maximal ATP production capacity and (**C**) predicted maximal glucose uptake capacity were similar between SSCs and progenitors. (**D**) Predicted maximal lactate uptake capacity was significantly higher in SSCs (paired t-test, p = 0.0457), suggesting potential differences in substrate utilisation between the two populations. Overall, the modelling indicates that metabolic differences between SSCs and progenitors likely reflect subtle shifts in pathway utilisation rather than large differences in total energy production capacity.

**Supplementary Table S1. Proteomic data for selected metabolic and mitochondrial proteins.**

**Supplementary Table S2. Quantitative TEM-derived mitochondrial and cristae parameters.**

**Supplementary Table S3. Descriptive and inferential statistics for Seahorse metabolic parameters in SSCs and progenitor spermatogonia.**

## REFERENCES

1. Acharya, S., E. Hanssen, V.B. Botha, T.M. Smith, S. Jayatissa, Z. Trifunovic, L.A. Bartho, J.E. Schjenken, T.u.J. Kaitu’u-Lino, A.V. Perkins, J.L. James, K.G. Pringle, J.C. Bouwer, R. Smith, and J.J. Fisher. 2025. Electron tomography reveals mitochondrial network and cristae remodelling during cell differentiation in the human placenta. bioRxiv: 2025.2006.2009.658204.

2. Acharya, S., E. Hanssen, V.B. Botha, T.M. Smith, S. Jayatissa, Z. Trifunovic, L.A. Bartho, J.E. Schjenken, T.u.J. Kaitu’u-Lino, A.V. Perkins, J.L. James, K.G. Pringle, J.C. Bouwer, R. Smith, and J.J. Fisher. 2026a. Electron tomography reveals mitochondrial network and cristae remodelling during cell differentiation in the human placenta. Nature Communications. 17:8407.

3. Acharya, S., E. Hanssen, J.C. Bouwer, J.E. Schjenken, K.G. Pringle, R. Smith, and J.J. Fisher. 2026b. Exploring placental ultrastructure: A review of electron microscopy techniques and emerging methods for resolving 3D organelle architecture. Placenta.

4. Afzal, N., W.J. Lederer, M.S. Jafri, and C.A. Mannella. 2021. Effect of crista morphology on mitochondrial ATP output: A computational study. Curr Res Physiol. 4:163–176.

5. Bahat, A., and A. Gross. 2019. Mitochondrial plasticity in cell fate regulation. J Biol Chem. 294:13852–13863.

6. Basu, K., D. Lajoie, T. Aumentado-Armstrong, J. Chen, R.I. Koning, B. Bossy, M. Bostina, A. Sik, E. Bossy-Wetzel, and I. Rouiller. 2017. Molecular mechanism of DRP1 assembly studied in vitro by cryo-electron microscopy. PloS one. 12:e0179397.

7. Belazi, D., S. Solé-Domènech, B. Johansson, M. Schalling, and P. Sjövall. 2009. Chemical analysis of osmium tetroxide staining in adipose tissue using imaging ToF-SIMS. Histochem Cell Biol. 132:105–115.

8. Berndt, N., J. Eckstein, I. Wallach, S. Nordmeyer, M. Kelm, M. Kirchner, L. Goubergrits, M. Schafstedde, A. Hennemuth, M. Kraus, T. Grune, P. Mertins, T. Kuehne, and H.G. Holzhütter. 2021. CARDIOKIN1: Computational Assessment of Myocardial Metabolic Capability in Healthy Controls and Patients With Valve Diseases. Circulation. 144:1926–1939.

9. Berndt, N., O. Kann, and H.G. Holzhütter. 2015. Physiology-based kinetic modeling of neuronal energy metabolism unravels the molecular basis of NAD(P)H fluorescence transients. J Cereb Blood Flow Metab. 35:1494–1506.

10. Bernstein, I.R., B. Nixon, J.M. Lyons, K.B. Damyanova, C.S. De Oliveira, N.S. Mabotuwana, S.J. Stanger, G.E. Kaiko, T.H. Ying, J.M. Oatley, N.M. Skillen, A.J. Lochrin, J.L. Peters, and T. Lord. 2023. The hypoxia inducible factor EPAS1 is required for spermatogonial stem cell function in regenerative conditions. iScience:108424.

11. Bibb, M.J., R.A. Van Etten, C.T. Wright, M.W. Walberg, and D.A. Clayton. 1981. Sequence and gene organization of mouse mitochondrial DNA. Cell. 26:167–180.

12. Bigarella, C.L., R. Liang, and S. Ghaffari. 2014. Stem cells and the impact of ROS signaling. Development. 141:4206–4218.

13. Boussouar, F., and M. Benahmed. 2004. Lactate and energy metabolism in male germ cells. Trends in Endocrinology & Metabolism. 15:345–350.

14. Buzzard, E., M. McLaren, P. Bragoszewski, A. Brancaccio, H. Ford, B. Daum, P. Kuwabara, I. Collinson, and V. Gold. 2024. The consequence of ATP synthase dimer angle on mitochondrial morphology studied by cryo-electron tomography. Biochem J. 481:161–175.

15. Chan, F., M.J. Oatley, A.V. Kaucher, Q.-E. Yang, C.J. Bieberich, C.S. Shashikant, and J.M. Oatley. 2014. Functional and molecular features of the Id4+ germline stem cell population in mouse testes. Genes & development. 28:1351–1362.

16. Chandel, N.S., H. Jasper, T.T. Ho, and E. Passegué. 2016. Metabolic regulation of stem cell function in tissue homeostasis and organismal ageing. Nature Cell Biology. 18:823–832.

17. Chen, W., Y. Sun, Q. Sun, J. Zhang, M. Jiang, C. Chang, X. Huang, C. Wang, P. Wang, Z. Zhang, X. Chen, and Y. Wang. 2020a. MFN2 Plays a Distinct Role from MFN1 in Regulating Spermatogonial Differentiation. Stem Cell Reports. 14:803–817.

18. Chen, W., Z. Zhang, C. Chang, Z. Yang, P. Wang, H. Fu, X. Wei, E. Chen, S. Tan, W. Huang, L. Sun, T. Ni, Y. Yang, and Y. Wang. 2020b. A bioenergetic shift is required for spermatogonial differentiation. Cell Discovery. 6:56.

19. Cho, Y.M., S. Kwon, Y.K. Pak, H.W. Seol, Y.M. Choi, D.J. Park, K.S. Park, and H.K. Lee. 2006. Dynamic changes in mitochondrial biogenesis and antioxidant enzymes during the spontaneous differentiation of human embryonic stem cells. Biochem Biophys Res Commun. 348:1472–1478.

20. Choi, J., B.J. Seo, H. La, S.H. Yoon, Y.J. Hong, J.H. Lee, H.M. Chung, K. Hong, and J.T. Do. 2020. Comparative analysis of the mitochondrial morphology, energy metabolism, and gene expression signatures in three types of blastocyst-derived stem cells. Redox Biol. 30:101437.

21. Cogliati, S., J.A. Enriquez, and L. Scorrano. 2016. Mitochondrial Cristae: Where Beauty Meets Functionality. Trends in Biochemical Sciences. 41:261–273.

22. Corcoran, C.C., C.R. Grady, T. Pisitkun, J. Parulekar, and M.A. Knepper. 2017. From 20th century metabolic wall charts to 21st century systems biology: database of mammalian metabolic enzymes. American Journal of Physiology-Renal Physiology. 312:F533–F542.

23. Čunátová, K., D.P. Reguera, J. Houštěk, T. Mráček, and P. Pecina. 2020. Role of cytochrome c oxidase nuclear-encoded subunits in health and disease. Physiol Res. 69:947–965.

24. Damyanova, K.B., B. Nixon, S.D. Johnston, A. Gambini, P.P. Benitez, and T. Lord. 2024. Spermatogonial stem cell technologies: applications from human medicine to wildlife conservation†. Biology of Reproduction.

25. De Oliveira, C.S., B. Nixon, and T. Lord. 2022. A scRNA-seq Approach to Identifying Changes in Spermatogonial Stem Cell Gene Expression Following in vitro Culture. Front Cell Dev Biol. 10:782996.

26. Facucho-Oliveira, J.M., J. Alderson, E.C. Spikings, S. Egginton, and J.C. St John. 2007. Mitochondrial DNA replication during differentiation of murine embryonic stem cells. J Cell Sci. 120:4025–4034.

27. Fukuda, R., H. Zhang, J.W. Kim, L. Shimoda, C.V. Dang, and G.L. Semenza. 2007. HIF-1 regulates cytochrome oxidase subunits to optimize efficiency of respiration in hypoxic cells. Cell. 129:111–122.

28. Gilkerson, R., P. De La Torre, and S. St. Vallier. 2021. Mitochondrial OMA1 and OPA1 as Gatekeepers of Organellar Structure/Function and Cellular Stress Response. Frontiers in Cell and Developmental Biology. Volume 9 – 2021.

29. Gottschalk, B., Z. Koshenov, M. Waldeck-Weiermair, S. Radulović, F.E. Oflaz, M. Hirtl, O.A. Bachkoenig, G. Leitinger, R. Malli, and W.F. Graier. 2022. MICU1 controls spatial membrane potential gradients and guides Ca(2+) fluxes within mitochondrial substructures. Commun Biol. 5:649.

30. Guarani, V., E.M. McNeill, J.A. Paulo, E.L. Huttlin, F. Fröhlich, S.P. Gygi, D. Van Vactor, and J.W. Harper. 2015. QIL1 is a novel mitochondrial protein required for MICOS complex stability and cristae morphology. Elife. 4.

31. Guo, J., E.J. Grow, C. Yi, H. Mlcochova, G.J. Maher, C. Lindskog, P.J. Murphy, C.L. Wike, D.T. Carrell, A. Goriely, J.M. Hotaling, and B.R. Cairns. 2017. Chromatin and Single-Cell RNA-Seq Profiling Reveal Dynamic Signaling and Metabolic Transitions during Human Spermatogonial Stem Cell Development. Cell Stem Cell. 21:533–546.e536.

32. Hahn, A., K. Parey, M. Bublitz, D.J. Mills, V. Zickermann, J. Vonck, W. Kühlbrandt, and T. Meier. 2016. Structure of a Complete ATP Synthase Dimer Reveals the Molecular Basis of Inner Mitochondrial Membrane Morphology. Mol Cell. 63:445–456.

33. Helsel, A.R., M.J. Oatley, and J.M. Oatley. 2017a. Glycolysis-Optimized Conditions Enhance Maintenance of Regenerative Integrity in Mouse Spermatogonial Stem Cells during Long-Term Culture. Stem Cell Reports. 8:1430–1441.

34. Helsel, A.R., Q. Yang, M.J. Oatley, T. Lord, F. Sablitzky, and J.M. Oatley. 2017b. ID4 levels dictate the stem cell state in mouse spermatogonia. Development. 144:624–634.

35. Hermann, B.P., K. Cheng, A. Singh, L. Roa-De La Cruz, K.N. Mutoji, I.C. Chen, H. Gildersleeve, J.D. Lehle, M. Mayo, B. Westernströer, N.C. Law, M.J. Oatley, E.K. Velte, B.A. Niedenberger, D. Fritze, S. Silber, C.B. Geyer, J.M. Oatley, and J.R. McCarrey. 2018. The Mammalian Spermatogenesis Single-Cell Transcriptome, from Spermatogonial Stem Cells to Spermatids. Cell Rep. 25:1650–1667.e1658.

36. Holmström, K.M., and T. Finkel. 2014. Cellular mechanisms and physiological consequences of redox-dependent signalling. Nature reviews. Molecular cell biology. 15:411–421.

37. Ito, K., and T. Suda. 2014. Metabolic requirements for the maintenance of self-renewing stem cells. Nat Rev Mol Cell Biol. 15:243–256.

38. Ito, M., M. Imai, M. Muraki, K. Miyado, J. Qin, S. Kyuwa, Y. Yoshikawa, Y. Hosoi, H. Saito, and Y. Takahashi. 2011. GSTT1 is upregulated by oxidative stress through p38-MK2 signaling pathway in human granulosa cells: possible association with mitochondrial activity. Aging (Albany NY). 3:1213–1223.

39. Javadirad, S.-M., and M. Mokhtari. 2021. TXNDC2 joint molecular marker is associated with testis pathology and is an accurate predictor of sperm retrieval. Scientific Reports. 11:13064.

40. Jensen, R.B., and E. Rothenberg. 2020. Preserving genome integrity in human cells via DNA double-strand break repair. Mol Biol Cell. 31:859–865.

41. Jiang, L., M. Wang, S. Lin, R. Jian, X. Li, J. Chan, G. Dong, H. Fang, A.E. Robinson, and M.P. Snyder. 2020. A Quantitative Proteome Map of the Human Body. Cell. 183:269–283.e219.

42. Kanatsu-Shinohara, M., T. Tanaka, N. Ogonuki, A. Ogura, H. Morimoto, P.F. Cheng, R.N. Eisenman, A. Trumpp, and T. Shinohara. 2016. Myc/Mycn-mediated glycolysis enhances mouse spermatogonial stem cell self-renewal. Genes & Development. 30:2637–2648.

43. Katti, P., P. Prasad, S.K. Masenga, P. Venkhatesh, Z. Vue, A.G. Marshall, B. Rodriguez, H. Le, E. Garza-Lopez, A. Murphy, B. Jenkins, A. Kadam, J. Shao, A. Crabtree, P. Martin, C. Evans, M.A. Phillips, D. Hubert, N. Wandira, O.M. Ochayi, D. Tomar, C.R. Williams, J. Gaddy, B. Tomeau, L. Bell, T. Gillyard, M. Hamilton, V. Sharma, M.M. Khan, E. Zaganjor, O.A. Ajijola, E. Scudese, T.W. Miller Fleming, A. Kinder, C. Dash, A.M. Quintana, B.C. Mobley, J.D. Berry, P. Jadiya, D.-F. Dai, A. Kirabo, O. Kovtun, J.C. Schafer, S. Schaffer, R.O. Pereira, M.R. McReynolds, and A. Hinton. 2026. The MICOS Complex Regulates Mitochondrial Structure and Oxidative Stress During Age-Dependent Structural Deficits in the Kidney. bioRxiv: 2024.2006.2009.598108.

44. Khacho, M., A. Clark, D.S. Svoboda, J. Azzi, J.G. MacLaurin, C. Meghaizel, H. Sesaki, D.C. Lagace, M. Germain, M.E. Harper, D.S. Park, and R.S. Slack. 2016. Mitochondrial Dynamics Impacts Stem Cell Identity and Fate Decisions by Regulating a Nuclear Transcriptional Program. Cell Stem Cell. 19:232–247.

45. Koirala, S., Q. Guo, R. Kalia, H.T. Bui, D.M. Eckert, A. Frost, and J.M. Shaw. 2013. Interchangeable adaptors regulate mitochondrial dynamin assembly for membrane scission. Proc Natl Acad Sci U S A. 110:E1342–1351.

46. Kondadi, A.K., R. Anand, and A.S. Reichert. 2020. Cristae Membrane Dynamics – A Paradigm Change. Trends Cell Biol. 30:923–936.

47. Kubota, H., M.R. Avarbock, J.A. Schmidt, and R.L. Brinster. 2009. Spermatogonial stem cells derived from infertile Wv/Wv mice self-renew in vitro and generate progeny following transplantation. Biol Reprod. 81:293–301.

48. La, H.M., J. Liao, J.M.D. Legrand, F.J. Rossello, A.L. Chan, V. Vaghjiani, J.E. Cain, A. Papa, T.L. Lee, and R.M. Hobbs. 2022. Distinctive molecular features of regenerative stem cells in the damaged male germline. Nat Commun. 13:2500.

49. Lam, J., P. Katti, M. Biete, M. Mungai, S. AshShareef, K. Neikirk, E. Garza Lopez, Z. Vue, T.A. Christensen, H.K. Beasley, T.A. Rodman, S.A. Murray, J.L. Salisbury, B. Glancy, J. Shao, R.O. Pereira, E.D. Abel, and A. Hinton, Jr. 2021. A Universal Approach to Analyzing Transmission Electron Microscopy with ImageJ. Cells. 10.

50. Li, T., Z. Tang, Y. Song, M. Su, X. Liu, F. Li, X. Wei, X. Luo, B. Zhou, Y. Wang, and L. Zhang. 2026. Busulfan plus cyclophosphamide induced spermatogenic dysfunction and recovery: A dynamic change perspective. Toxicology Letters. 417:111833.

51. Lillig, C.H., C. Berndt, O. Vergnolle, M.E. Lönn, C. Hudemann, E. Bill, and A. Holmgren. 2005. Characterization of human glutaredoxin 2 as iron–sulfur protein: A possible role as redox sensor. Proceedings of the National Academy of Sciences. 102:8168–8173.

52. Liu, X., Q. Li, W. Wang, and F. Liu. 2019. Aberrant expression of sperm-specific glycolytic enzymes are associated with poor sperm quality. Mol Med Rep. 19:2471–2478.

53. Lord, T., and B. Nixon. 2020. Metabolic Changes Accompanying Spermatogonial Stem Cell Differentiation. Developmental Cell. 52:399–411.

54. Lord, T., and J.M. Oatley. 2017. Regulation of Spermatogonial Stem Cell Maintenance and Self-Renewal. *In* The Biology of Mammalian Spermatogonia. J.M. Oatley and M.D. Griswold, editors. Springer New York, New York, NY. 91–129.

55. Losón, O.C., Z. Song, H. Chen, and D.C. Chan. 2013. Fis1, Mff, MiD49, and MiD51 mediate Drp1 recruitment in mitochondrial fission. Mol Biol Cell. 24:659–667.

56. Mannella, C.A. 2006. Structure and dynamics of the mitochondrial inner membrane cristae. Biochimica et Biophysica Acta (BBA) – Molecular Cell Research. 1763:542–548.

57. Martin, L.A., and M. Seandel. 2013. Serial enrichment of spermatogonial stem and progenitor cells (SSCs) in culture for derivation of long-term adult mouse SSC lines. J Vis Exp: e50017.

58. Mishmar, D., E. Ruiz-Pesini, M. Brandon, and D.C. Wallace. 2004. Mitochondrial DNA-like sequences in the nucleus (NUMTs): insights into our African origins and the mechanism of foreign DNA integration. Hum Mutat. 23:125–133.

59. Mori, Y., N. Ogonuki, A. Hasegawa, M. Kanatsu-Shinohara, A. Ogura, Y. Wang, J.R. McCarrey, and T. Shinohara. 2021. OGG1 protects mouse spermatogonial stem cells from reactive oxygen species in culture†. Biol Reprod. 104:706–716.

60. Morimoto, H., K. Iwata, N. Ogonuki, K. Inoue, O. Atsuo, M. Kanatsu-Shinohara, T. Morimoto, C. Yabe-Nishimura, and T. Shinohara. 2013. ROS are required for mouse spermatogonial stem cell self-renewal. Cell stem cell. 12:774–786.

61. Morimoto, H., T. Yamamoto, T. Miyazaki, N. Ogonuki, A. Ogura, T. Tanaka, M. Kanatsu-Shinohara, C. Yabe-Nishimura, H. Zhang, Y. Pommier, A. Trumpp, and T. Shinohara. 2021. An interplay of NOX1-derived ROS and oxygen determines the spermatogonial stem cell self-renewal efficiency under hypoxia. Genes Dev. 35:250–260.

62. Nolden, K.A., M.C. Harwig, and R.B. Hill. 2023. Human Fis1 directly interacts with Drp1 in an evolutionarily conserved manner to promote mitochondrial fission. J Biol Chem. 299:105380.

63. Oatley, J.M., and R.L. Brinster. 2012. The germline stem cell niche unit in mammalian testes. Physiol Rev. 92:577–595.

64. Perkins, G., C. Renken, M.E. Martone, S.J. Young, M. Ellisman, and T. Frey. 1997. Electron tomography of neuronal mitochondria: three-dimensional structure and organization of cristae and membrane contacts. J Struct Biol. 119:260–272.

65. Pham, L., T. Arroum, J. Wan, L. Pavelich, J. Bell, P.T. Morse, I. Lee, L.I. Grossman, T.H. Sanderson, M.H. Malek, and M. Hüttemann. 2024. Regulation of mitochondrial oxidative phosphorylation through tight control of cytochrome c oxidase in health and disease – Implications for ischemia/reperfusion injury, inflammatory diseases, diabetes, and cancer. Redox Biology. 78:103426.

66. Picard, M., K. White, and D.M. Turnbull. 2013. Mitochondrial morphology, topology, and membrane interactions in skeletal muscle: a quantitative three-dimensional electron microscopy study. J Appl Physiol (1985). 114:161–171.

67. Rampazzo, C., L. Gallinaro, E. Milanesi, E. Frigimelica, P. Reichard, and V. Bianchi. 2000. A deoxyribonucleotidase in mitochondria: involvement in regulation of dNTP pools and possible link to genetic disease. Proc Natl Acad Sci U S A. 97:8239–8244.

68. Rath, S., R. Sharma, R. Gupta, T. Ast, C. Chan, T.J. Durham, R.P. Goodman, Z. Grabarek, M.E. Haas, W.H.W. Hung, P.R. Joshi, A.A. Jourdain, S.H. Kim, A.V. Kotrys, S.S. Lam, J.G. McCoy, J.D. Meisel, M. Miranda, A. Panda, A. Patgiri, R. Rogers, S. Sadre, H. Shah, O.S. Skinner, T.-L. To, Melissa A. Walker, H. Wang, P.S. Ward, J. Wengrod, C.-C. Yuan, S.E. Calvo, and V.K. Mootha. 2020. MitoCarta3.0: an updated mitochondrial proteome now with sub-organelle localization and pathway annotations. Nucleic Acids Research. 49:D1541–D1547.

69. Rydström, J. 2006. Mitochondrial NADPH, transhydrogenase and disease. Biochim Biophys Acta. 1757:721–726.

70. Schmidt, J.A., L.K. Abramowitz, H. Kubota, X. Wu, Z. Niu, M.R. Avarbock, J.W. Tobias, M.S. Bartolomei, and R.L. Brinster. 2010. In Vivo and In Vitro Aging Is Detrimental to Mouse Spermatogonial Stem Cell Function1. Biology of Reproduction. 84:698–706.

71. Skerrett-Byrne, D.A., B. Nixon, A. Sanz-Moreno, F. Dámek, P. da Silva-Buttkus, R. Teperino, V. Gailus-Durner, H. Fuchs, M. Hrabé de Angelis, J.M. Oatley, C. Cason, I.R. Bernstein, and T. Lord. 2026. A proteomic and phosphoproteomic comparison of mouse spermatogonial stem cells and progenitor spermatogonia. bioRxiv: 2026.2007.2015.738804.

72. Smith, T.B., M.A. Baker, H.S. Connaughton, U. Habenicht, and R.J. Aitken. 2013. Functional deletion of Txndc2 and Txndc3 increases the susceptibility of spermatozoa to age-related oxidative stress. Free Radical Biology and Medicine. 65:872–881.

73. Stachowiak, J.C., F.M. Brodsky, and E.A. Miller. 2013. A cost-benefit analysis of the physical mechanisms of membrane curvature. Nat Cell Biol. 15:1019–1027.

74. Strauss, M., G. Hofhaus, R.R. Schröder, and W. Kühlbrandt. 2008. Dimer ribbons of ATP synthase shape the inner mitochondrial membrane. Embo j. 27:1154–1160.

75. Sukonina, V., H. Ma, W. Zhang, S. Bartesaghi, S. Subhash, M. Heglind, H. Foyn, M.J. Betz, D. Nilsson, M.E. Lidell, J. Naumann, S. Haufs-Brusberg, H. Palmgren, T. Mondal, M. Beg, M.P. Jedrychowski, K. Taskén, A. Pfeifer, X.-R. Peng, C. Kanduri, and S. Enerbäck. 2019. FOXK1 and FOXK2 regulate aerobic glycolysis. Nature. 566:279–283.

76. Sundaresan, N.R., M. Gupta, G. Kim, S.B. Rajamohan, A. Isbatan, and M.P. Gupta. 2009. Sirt3 blocks the cardiac hypertrophic response by augmenting Foxo3a-dependent antioxidant defense mechanisms in mice. J Clin Invest. 119:2758–2771.

77. Tagelenbosch, R.A.J., and D.G. de Rooij. 1993. A quantitative study of spermatogonial multiplication and stem cell renewal in the C3H/101 F1 hybrid mouse. Mutation Research/Fundamental and Molecular Mechanisms of Mutagenesis. 290:193–200.

78. Tang, A.H., and T.A. Rando. 2014. Induction of autophagy supports the bioenergetic demands of quiescent muscle stem cell activation. The EMBO journal. 33:2782–2797.

79. TeSlaa, T., M. Ralser, J. Fan, and J.D. Rabinowitz. 2023. The pentose phosphate pathway in health and disease. Nature Metabolism. 5:1275–1289.

80. Tomar, D., M. Thomas, J.F. Garbincius, D.W. Kolmetzky, O. Salik, P. Jadiya, S.K. Joseph, A.C. Carpenter, G. Hajnóczky, and J.W. Elrod. 2023. MICU1 regulates mitochondrial cristae structure and function independently of the mitochondrial Ca(2+) uniporter channel. Sci Signal. 16:eabi8948.

81. Urish, K.L., J.B. Vella, M. Okada, B.M. Deasy, K. Tobita, B.B. Keller, B. Cao, J.D. Piganelli, and J. Huard. 2009. Antioxidant levels represent a major determinant in the regenerative capacity of muscle stem cells. Mol Biol Cell. 20:509–520.

82. Ushio-Fukai, M., and J. Rehman. 2014. Redox and metabolic regulation of stem/progenitor cells and their niche. Antioxid Redox Signal. 21:1587–1590.

83. Varum, S., A.S. Rodrigues, M.B. Moura, O. Momcilovic, C.A.t. Easley, J. Ramalho-Santos, B. Van Houten, and G. Schatten. 2011. Energy metabolism in human pluripotent stem cells and their differentiated counterparts. PloS one. 6:e20914.

84. Varuzhanyan, G., and D.C. Chan. 2020. Mitochondrial dynamics during spermatogenesis. J Cell Sci. 133.

85. Varuzhanyan, G., R. Rojansky, M.J. Sweredoski, R.L.J. Graham, S. Hess, M.S. Ladinsky, and D.C. Chan. 2019. Mitochondrial fusion is required for spermatogonial differentiation and meiosis. Elife. 8.

86. Vesell, E.S. 1965. Lactate dehydrogenase isozymes: substrate inhibition in various human tissues. *Science (New York*, N.Y.). 150:1590–1593.

87. Voigt, A.L., R. Dardari, L. Su, N.L.M. Lara, S. Sinha, A. Jaffer, S.K. Munyoki, W. Alpaugh, A. Dufour, J. Biernaskie, K.E. Orwig, and I. Dobrinski. 2022. Metabolic transitions define spermatogonial stem cell maturation. Human Reproduction. 37:2095–2112.

88. Wanet, A., T. Arnould, M. Najimi, and P. Renard. 2015. Connecting Mitochondria, Metabolism, and Stem Cell Fate. Stem Cells Dev. 24:1957–1971.

89. Westrate, L.M., J.A. Drocco, K.R. Martin, W.S. Hlavacek, and J.P. MacKeigan. 2014. Mitochondrial Morphological Features Are Associated with Fission and Fusion Events. PloS one. 9:e95265.

90. Xi, H.-M., Y.-J. Ren, F. Ren, Y. Li, T.-Y. Feng, Z. Wang, Y.-Q. Du, L.-K. Zhang, and J.-H. Hu. 2022. Recent advances in isolation, identification, and culture of mammalian spermatogonial stem cells. Asian Journal of Andrology. 24:5–14.

91. Zhang, Z., J. Miao, H. Wang, I. Ali, D. Nguyen, W. Chen, and Y. Wang. 2024. Accelerated mitochondrial dynamics promote spermatogonial differentiation. Stem Cell Reports.

