## Supplemental Figures for "Mitochondrial and metabolic plasticity distinguish stem cells from progenitors in the male germline"

### FIGURE S1

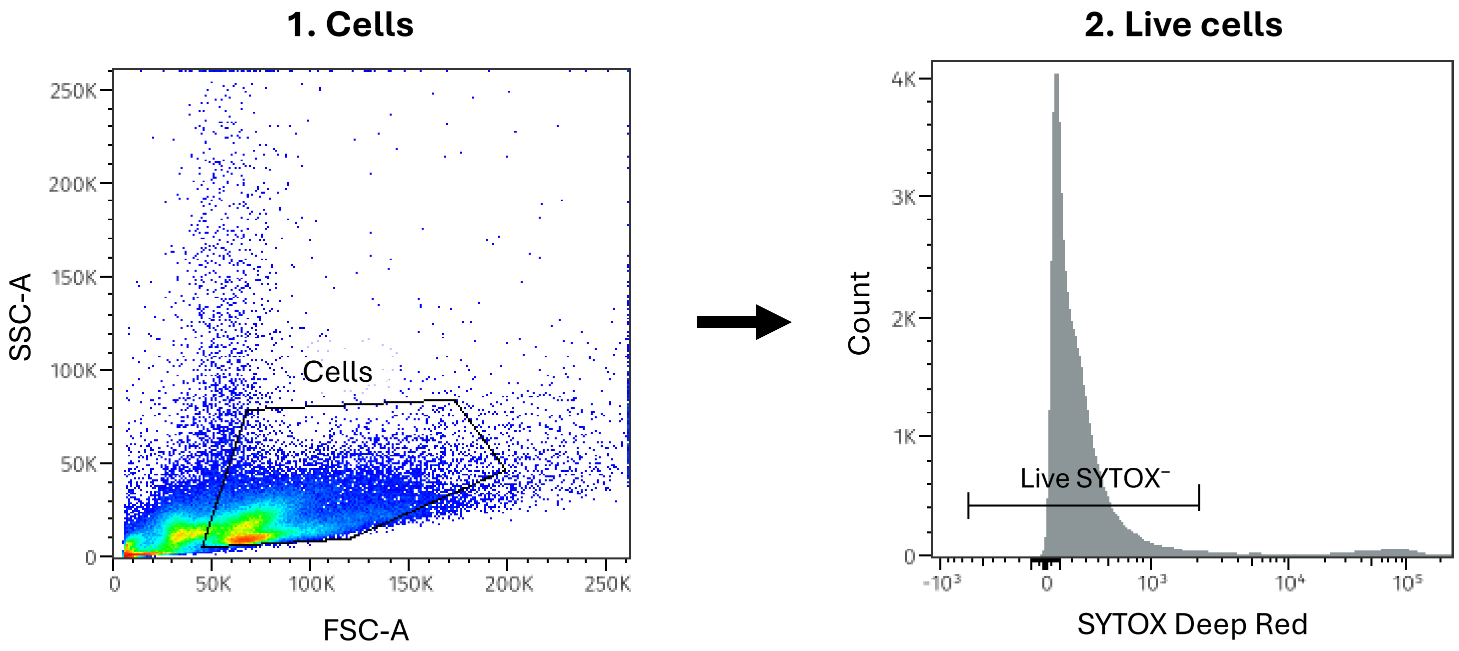


**Supplementary Figure S1. Representative gating strategy for MitoSOX Red flow cytometry analysis.**

Sequential gating strategy applied to testicular cell suspensions prior to identification of ID4-eGFP subpopulations. Cells were first gated on FSC-A/SSC-A to exclude debris and non-cellular events (**1**, "Cells"), followed by exclusion of non-viable cells based on SYTOX Deep Red fluorescence (**2**, "Live cells"; SYTOX$^{-}$ gate indicated). Downstream identification of ID4-eGFPBright and ID4-eGFPDim subpopulations and MitoSOX Red-positive gating is shown in **Fig. 7C**.

### FIGURE S2

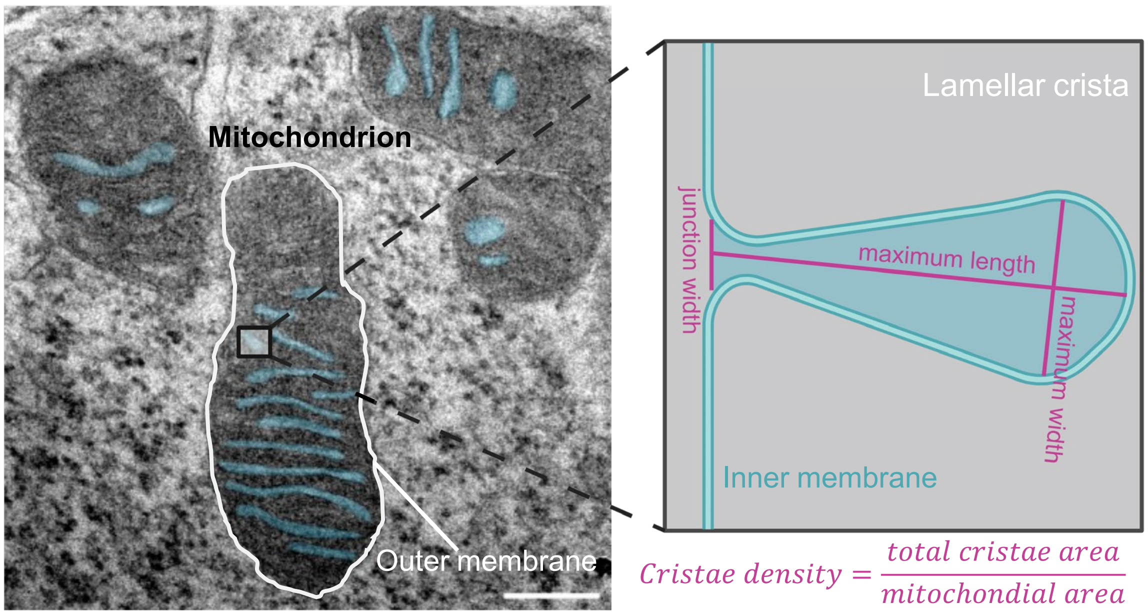


**Supplementary Figure S2. Definition of lamellar cristae measurements derived from 2D TEM images.**

Representative transmission electron micrograph of a spermatogonial mitochondrion illustrating lamellar cristae (left), with cristae manually segmented (blue) for quantitative analysis. The inset schematic (right) depicts the parameters used to quantify lamellar cristae morphology from 2D TEM sections, including maximum crista length, maximum crista width, and crista junction width at the point of connection to the inner mitochondrial membrane. Cristae density was calculated as the ratio of total cristae surface area to mitochondrial surface area. Measurements were performed only on clearly discernible lamellar cristae with well-defined membrane boundaries. Quantification was conducted in two dimensions and reflects the apparent cristae geometry within individual TEM sections. Scale bar = 200nm. Schematic is not to scale.

**FIGURE S3**

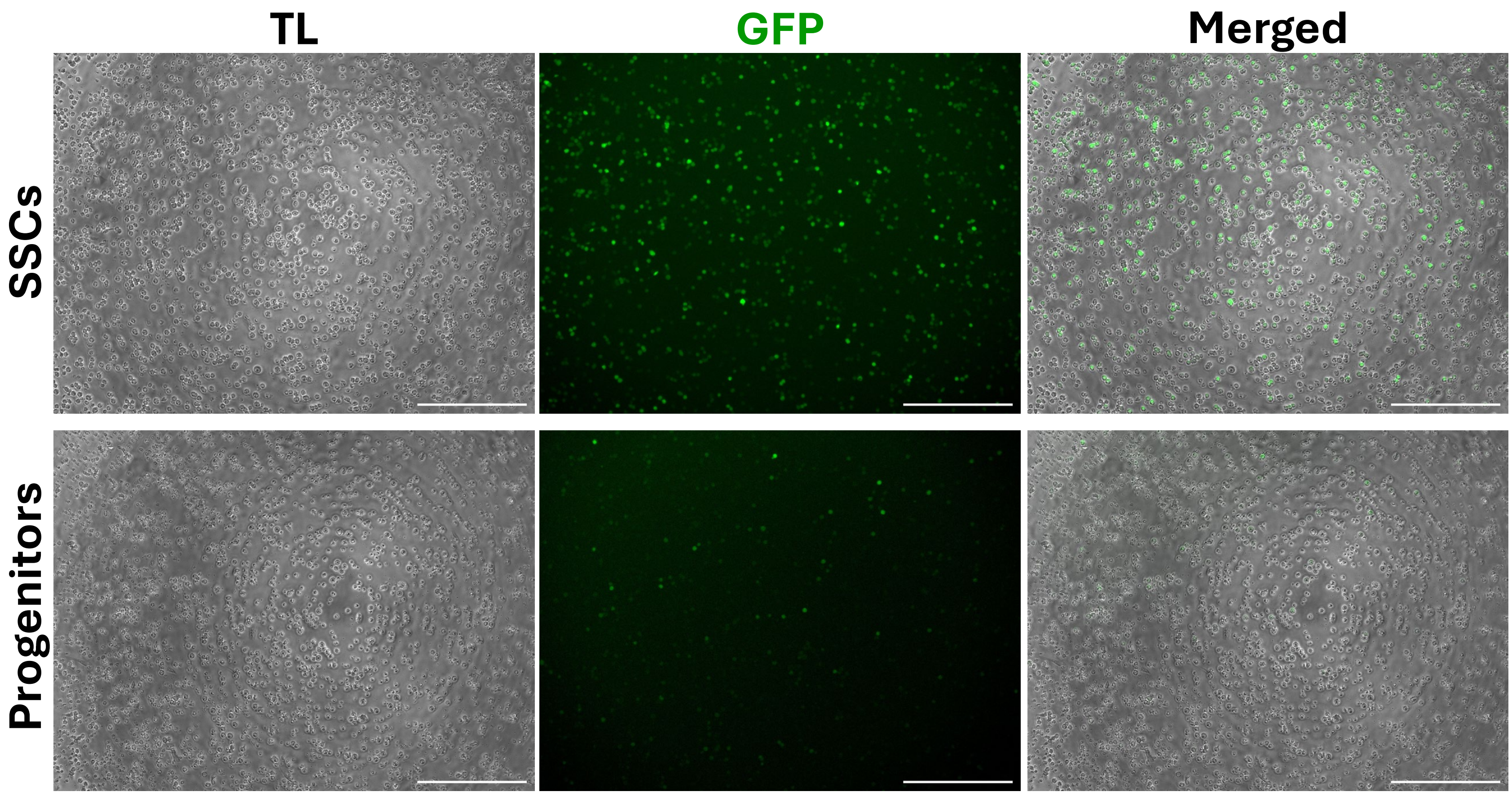


**Supplementary Figure S3.**
Representative images of ID4-eGFP$^{Bright}$ or ID4-eGFP$^{Dim}$ cells seeded in XF96 cell culture microplates at 50,000 cells/well for Seahorse metabolic assays after an overnight incubation at 37°C in 10% $O_2$ and 5% $CO_2$. Scale bar = 300µm.

### FIGURE S4

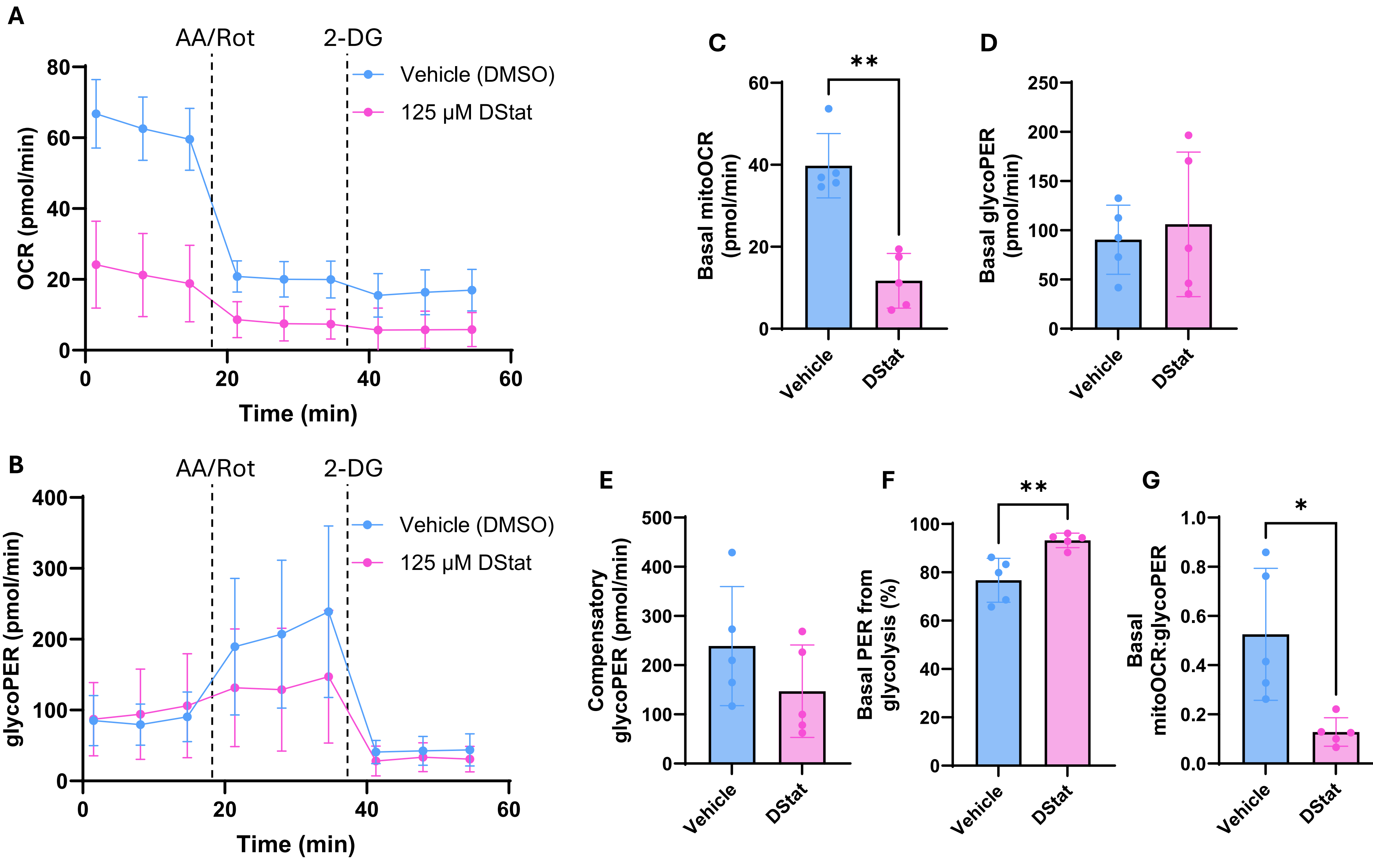


**Supplementary Figure S4. Daprodustat suppresses mitochondrial respiration and shifts cultured spermatogonia toward glycolysis.**

Primary cultures of undifferentiated spermatogonia were treated with 125 µM Daprodustat (DStat) or DMSO vehicle for 24 h at 10% $O_2$, before the Seahorse Glycolytic Rate Assay at 21% $O_2$. (**A–B**) Kinetic traces of oxygen consumption rate (OCR; **A**) and glycolytic proton efflux rate (glycoPER; **B**), shown as mean ± SD. Dashed lines indicate the injections of antimycin A/rotenone (AA/Rot) and 2-deoxy-D-glucose (2-DG). (**C–G**) Basal mitochondrial OCR (mitoOCR; C), basal glycoPER (**D**), compensatory glycoPER (**E**), percentage of basal PER derived from glycolysis (**F**), and ratio of basal mitoOCR to glycoPER (**G**). Bars show mean ± SD, and each point is one biological replicate (an independent culture; mean of 2–3 technical wells). Statistical comparisons were performed using paired two-tailed t-tests (n = 5). $^*p < 0.05$, $^{**}p < 0.01$.

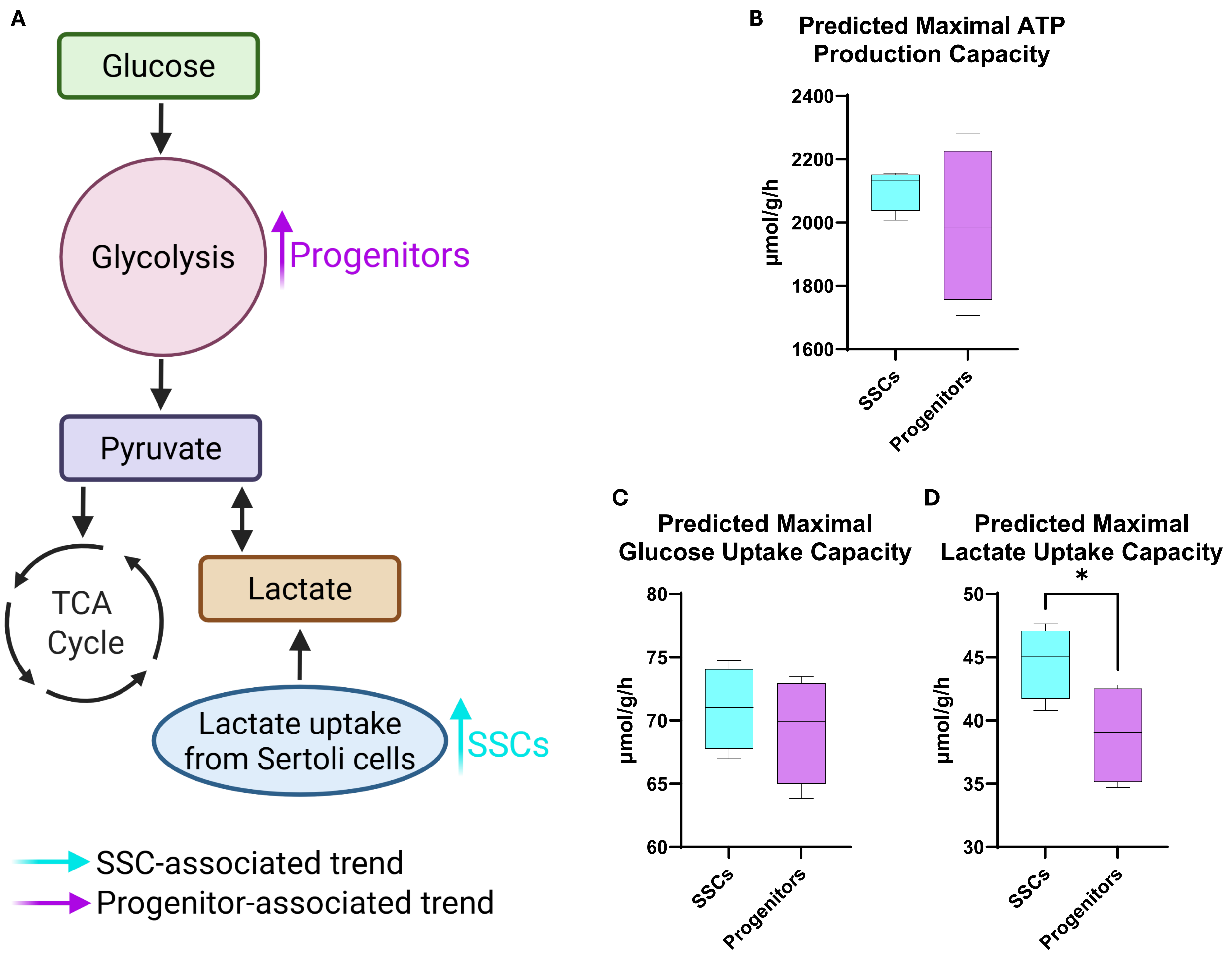


**Supplementary Figure S5. Quantitative systems metabolism (QSM) modelling of central carbon metabolism in SSCs and progenitor spermatogonia.** (**A**) Simplified schematic summarising predicted metabolic trends inferred from proteomic data. Progenitor spermatogonia show relatively greater representation of glycolytic enzymes, whereas SSCs display a tendency toward increased lactate uptake from the seminiferous microenvironment. (**B–D**) QSM modelling outputs showing predicted maximal ATP production capacity and substrate uptake capacities derived from protein abundance values. (**B**) Predicted maximal ATP production capacity and (**C**) predicted maximal glucose uptake capacity were similar between SSCs and progenitors. (**D**) Predicted maximal lactate uptake capacity was significantly higher in SSCs (paired t-test, $p = 0.0457$), suggesting potential differences in substrate utilisation between the two populations. Overall, the modelling indicates that metabolic differences between SSCs and progenitors likely reflect subtle shifts in pathway utilisation rather than large differences in total energy production capacity.
